# Native β-barrel substrates pass through two shared intermediates during folding on the BAM complex

**DOI:** 10.1101/2024.05.09.593409

**Authors:** Thiago M. A. Santos, Benjamin D. Thomson, Melissa D. Marquez, Lydia Pan, Tabasom H Monfared, Daniel E. Kahne

## Abstract

The assembly of β-barrel proteins into membranes is mediated by the evolutionarily conserved BAM complex. In *Escherichia coli*, BAM folds numerous substrates which vary considerably in size and shape. How BAM is able to efficiently fold such a diverse array of β-barrel substrates is not clear. Here, we develop a disulfide crosslinking method to trap native substrates in vivo as they fold on BAM. By placing a cysteine within the luminal wall of the BamA barrel as well as in the substrate β-strands, we can compare the residence time of each substrate strand within the BamA lumen. We validated this method using two defective, slow-folding substrates. We used this method to characterize stable intermediates which occur during folding of two structurally different native substrates. Strikingly, these intermediates occur during identical stages of folding for both substrates: soon after folding has begun, and just before folding is completed. We suggest that these intermediates arise due to barriers to folding that are common between β-barrel substrates, and that the BAM catalyst is able to fold so many different substrates because it addresses these common challenges.

**Significance Statement:** The outer membrane of Gram-negative bacteria is a barrier which protects these organisms from many antimicrobial agents. Here, we study the machine responsible for folding and inserting integral β-barrel proteins into the membrane: BAM. Outer membrane integrity and cell viability is dependent on the proper function of BAM. Here we show that stable intermediates exist on the folding pathway of native substrates. We also show that mutant substrates that increase the stability of these native intermediates can stall during folding. This creates permeability defects that can be exploited by antibiotics that normally do not cross the outer membrane. These observations could enable the design of strategies to combat Gram-negative pathogens.

## Introduction

Transmembrane proteins are composed of bundles of α-helices or a cylindrical array of antiparallel β-strands that forms a β-barrel. Both structural classes require conserved machines to facilitate their folding and assembly (1, 2). The bacterial β-<u>b</u>arrel <u>a</u>ssembly <u>m</u>achine (BAM) contains five conserved proteins: BamA, which itself is a 16-stranded β-barrel protein (Figure 1A and 1B), and the four associated lipoproteins BamBCDE (3–5). Of these five proteins, BamA and BamD are essential (3, 4). Unlike most enzymes, which act upon one or a small number of substrates, the BAM complex can accelerate the folding of numerous different β-barrel substrates of varying size (6–11). Our hypothesis is that, because the BAM complex can function as a general catalyst to fold so many different substrates, the folding pathways of these diverse substrates must share common features.

**Figure 1:**
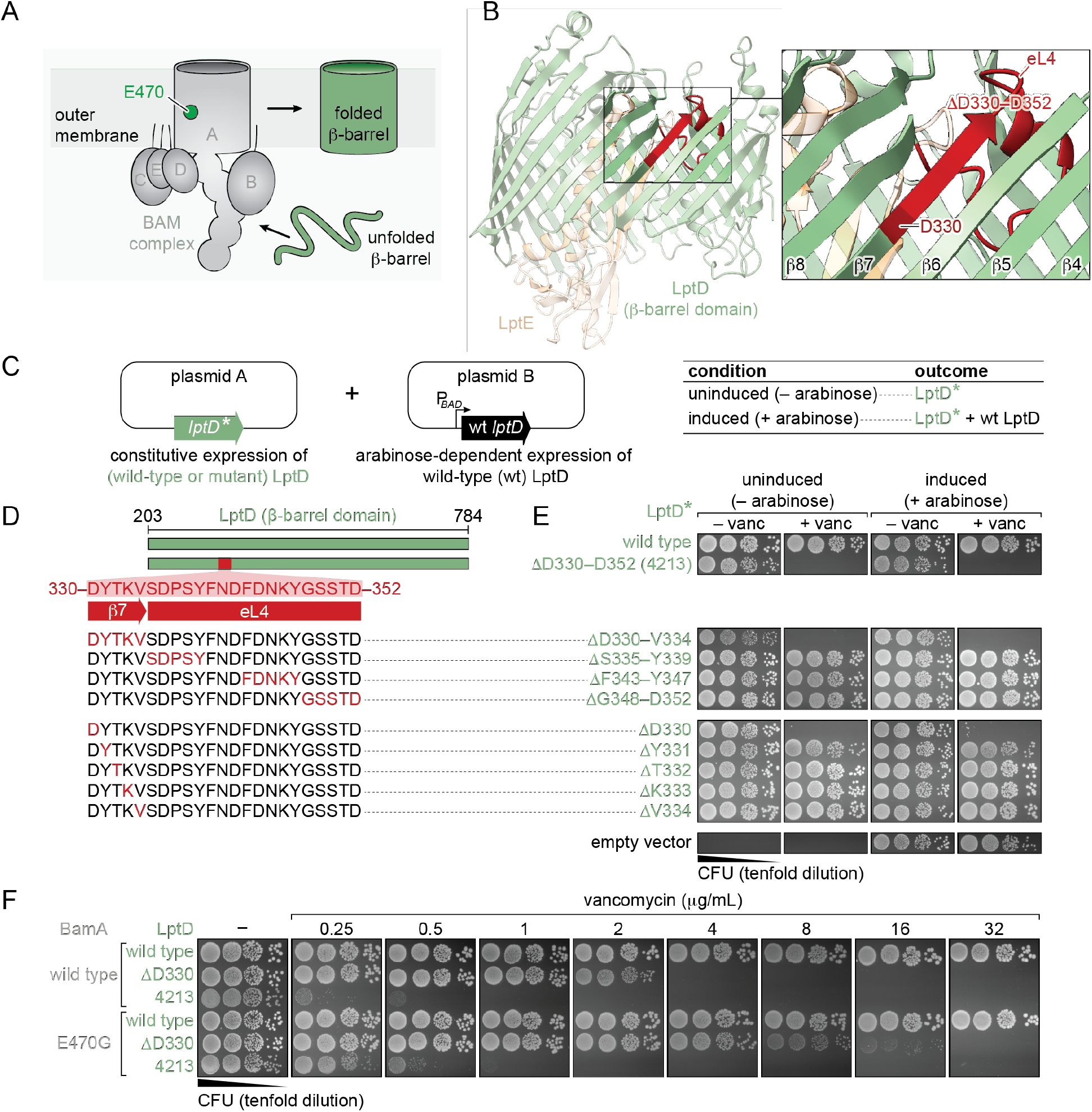
LptD-ΔD330 is the minimal deletion required to cause LptD4213 assembly defects on BAM. (A) BAM catalyzes the folding of outer membrane β-barrels. E470 lies on the interior surface of the BamA lumen. (B) β-barrel domain of LptD (PDB: 4RHB). LptD4213 is a slow-folding variant of LptD due to the deletion of residues D330-D352 (highlighted in red). D330 (dark red) is the N-terminal-most residue within the deleted region. (C) Two-plasmid system used to probe cell permeability defects caused by deletions within LptD. (D) Deleted regions tested (red) span the LptD4213 23-amino-acid deletion. (E) Plating assay of *E. coli* containing plasmids A and B on plates with and without arabinose or vancomycin. LptD-ΔD330 recapitulates the permeability defects caused by LptD4213. (F) Vancomycin sensitivity assay of *E. coli* containing LptD-ΔD330 and LptD4213 in cells with or without a chromosomal copy of BamA-E470G. Both LptD-ΔD330 and LptD4213 are partially rescued by BamA-E470G.

The BAM complex is an attractive potential target for antibiotics (12–17). Identifying the conserved features of BAM-catalyzed folding of multiple different substrates could inform what parts of the folding pathway can be targeted with antibiotics. In previous work (11, 18–20), we have suggested the following model for BAM-mediated folding of β-barrels (Figure S1A): substrates first interact with the BAM complex by binding to BamD via the conserved β-signal near the substrate C-terminus (18). The C-terminus of the substrate then binds to the N-terminal edge of the BamA barrel, creating a machine-substrate hybrid barrel (11). As folding proceeds, additional substrate strands are added to this hybrid barrel within the lumen of machine BamA and the folded substrate bulges into the membrane. Substrate release occurs after the entire substrate barrel has been folded, and the backbone hydrogen bonds that hold the machine N-terminus to the substrate C-terminus are exchanged pairwise in a zipping motion (20, 21). While this model contains a description of the landmarks that could be observed during folding, the information that has led to this model, and to alternative models (22–28), has been acquired primarily by studying substrates that have been structurally altered to slow down or prevent completion of their folding. It is not known if stable intermediates normally exist during the folding of native substrates of the BAM complex.

Of the several β-barrel substrates in *Escherichia coli*, we have selected the two that are essential for survival: BamA as a substrate (hereafter BamA^S^) and LptD. BamA within an active BAM complex (hereafter BamA^M^) folds other copies of BamA^S^ (20, 29), which interact with the lipoproteins BamBCDE to become a functional BamA^M^. LptD and the lipoprotein LptE comprise the outer membrane translocon required to transport lipopolysaccharide from the periplasm to the surface of the cell (30–32) (Figure 1C). The outer membrane lipoprotein LptE must associate with the BAM complex so that LptD can fold around it (33). This added complexity to fold LptD and the large size (26 β-strands) of LptD are related to why this substrate folds much more slowly on the BAM complex than any other barrel (9, 11, 33).

We have previously characterized two slow-folding mutants of LptD (11, 33). One of these two mutants, LptD4213, contains the 23 amino acid deletion ΔD330–D352 (34). It is missing part of β-strand 7 and nearly the entire extracellular loop 4 (Figures 1C [*segment highlighted in red*], S1B, and S1. The other mutant substrate, LptD-ΔD330, lacks only the aspartate residue at position 330 (Figure 1C) (11).

In this work, we continued to study these two slow-folding LptD mutant substrates. We establish that LptD-ΔD330, the minimalist deletion of LptD4213, exhibits the same fundamental folding defect as LptD4213. To gain this insight, we developed a method to detect stable interaction between regions of these slow-folding substrates and the luminal wall of the machine BamA. Using these tools, we identified two common intermediates that occur along the folding pathway for each of the slow-folding substrates LptD4213 and LptD-Δ D330. We also examined the effect of two suppressors of the folding defects imparted by the amino acid deletions in the slow-folding substrates, and we used the suppressors to validate that the intermediate states in BAM-catalyzed folding of both slow-folding substrates are on-pathway. Importantly, we found that these same intermediates exist during folding of wild-type LptD and BamA^S^ on the BAM complex as well. We argue that the folding intermediates we characterize here are conserved in different native substrates.

## Results

### Deletion of a single amino acid in LptD-ΔD330 produces similar phenotypes to the 23-amino acid deletion present in LptD4213

LptD4213 contains a deletion of 23 amino acids and stalls on the BAM complex during folding (11, 33). LptD-Δ D330, in which only the N-terminal most residue within the 23-amino acid deletion is missing, is also slow to fold (11). To determine which parts of the 23-amino acid region are linked to the characteristic phenotypes of LptD4213 in vivo, we followed a previously reported procedure to assess the permeability barrier of the outer membrane of *E. coli* cells expressing mutant variants of LptD (11, 19, 35). Cells expressing LptD4213 display increased sensitivity to antibiotics, which can be used as a proxy for outer membrane permeability. We constructed a series of plasmids (plasmid A) that constitutively express either a copy of wild-type LptD, or variants of LptD harboring deletions within the 23-amino acid region that defines LptD4213. We then transformed cells lacking their native chromosomal copy of the gene encoding LptD but harboring a second plasmid (plasmid B) carrying an arabinose-inducible copy of the gene encoding wild-type LptD (Figures 1D and 1E). We plated cells containing both plasmids in the absence or in the presence of the antibiotic vancomycin. As previously established, when wild-type LptD is constitutively expressed (plasmid A), cells grow equally well in the presence or absence of vancomycin, regardless of whether arabinose is added to the media (Figures 1F). Additionally, cells containing constitutively expressed LptD4213 do not grow in plates containing vancomycin, even when a wild-type copy of LptD is induced with arabinose (plasmid B). Thus, LptD4213 makes *E. coli* susceptible to vancomycin, regardless of whether an additional wild-type copy of LptD is present in the cell. This has been interpreted to mean that the membrane permeability of cells containing LptD4213 comes from a defect in assembly of LptD4213 by the BAM complex rather than a defect in the function of the assembled LptD4213.

We used this experimental design to evaluate what portion(s) of the 23-amino-acid deletion is responsible for the slowed folding on the BAM complex. We prepared sequential five amino acid truncations spanning the region deleted in the mutant LptD4213. We reasoned that if the deletion of any of these regions interferes with the ability of the BAM complex to properly fold these substrates, they should cause outer membrane defects comparable to the defects observed in the parent mutant LptD4213. We found that only deletion of the first five amino acids (ΔD330–V334) prevented proper folding of the substrate on the BAM complex and resulted in the same dominant defect in outer membrane permeability (Figures 1E and 1F). We concluded that an important region for proper folding of LptD by the BAM complex involves a feature contained in the first five amino acids of the region deleted in the mutant LptD4213. We then deleted individual amino acids in this region and found that only ΔD330 resulted in the same dominant sensitivity to vancomycin (Figure 1F). Additional experiments (Figures S1D and S1E) suggest that changes in registry of β-strand 7, resulting from deleting a single amino acid, do not explain the folding defect observed. Additionally, deletion of loop 2, 4, or 8 of LptD were found to confer folding defects (Figure S1F). Evidently, certain positive and negative charges must make specific contacts that are important for folding or barrel stability. We concluded that LptD-ΔD330, the minimalist version of the deletion in LptD4213, exhibits the same fundamental folding defect as LptD4213.

The suppressor BamA-E470G restores the folding rate of LptD-ΔD330 to wild-type level. If the 23-amino acid deletion in LptD4213 disrupts folding of the substrate in a similar way to LptD-ΔD330, BamA-E470G should also alleviate some of the folding defects of LptD4213. We constructed strains harboring a chromosomal copy of the gene encoding BamA-E470G and either wild-type LptD, LptD-ΔD330, or LptD4213. We plated these engineered strains on media containing increasing concentrations of vancomycin and found that BamA-E470G decreases the sensitivity to vancomycin in both mutants (Figure 1G). While the absolute amount of vancomycin required to inhibit growth differs between the two mutants, the relative effect of BamA-E470G is comparable. These results, in conjunction with our previous studies showing that this suppressor of rescues the rate of folding of LptD-ΔD330 (11), suggest that the region of BamA near the residue E470 must plays an important role in folding of substrates.

### Slow-folding substrates accumulate near residue E470 of BamA

If the region of BamA containing the residue E470 is directly involved in folding of substrates, both folding-defective LptD substrates should accumulate proximal to E470. To test this hypothesis, we performed an in vivo photocrosslinking experiment (Figure 2A). Upon photoactivation by UV light, *p*Bpa forms covalent bonds to proximal moieties (Figure 2A) and the resulting high-molecular weight adducts can then be detected by immunoblotting. We had previously used this approach to examine the accumulation of LptD4213 on BamA (11) and identified nine residues in the β-barrel domain of BamA (β-strands 2–10), mostly along the interior surface, that showed UV-dependent crosslinking to this slow-folding substrate (Figure 2B, *yellow and blue labeled residues*). We repeated this experiment with LptD-ΔD330 and found that only two of these nine residues, S439 and Y468, of BamA crosslink to this slow-folding substrate (Figures 2B and 2C, *yellow labeled residues*). Residue S439 is on β-strand 2 near the lateral gate of BamA, which is a critical region of contact for substrates during folding (11, 20) (Figure 2B). Formation of a substrate-machine hybrid barrel near BamA S439 is a landmark that our mechanism predicts should be shared between many substrates of the BAM complex. Residue Y468, interestingly, faces the lumen adjacent to E470 on β-strand 4 of BamA. This observation is consistent with the hypothesis that the luminal wall region of BamA near E470 is a critical point of interaction between machine and substrate during folding.

**Figure 2:**
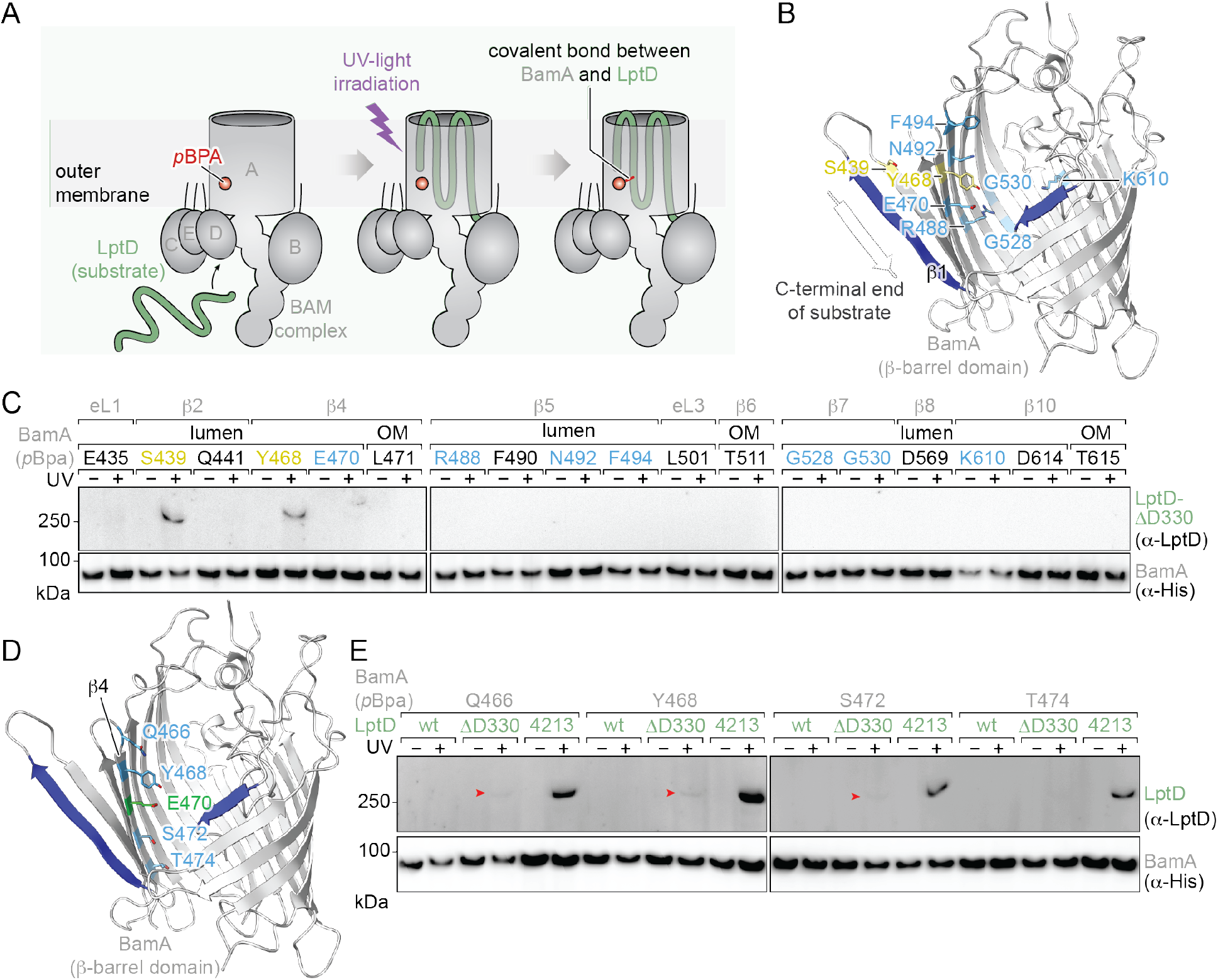
LptD-ΔD330 and LptD4213 accumulate near position E470 of BamA during folding. (A) In vivo photocrosslinking assay used to assess residence time of LptD variants within the BamA lumen. (B) Positions within BamA which crosslink to LptD4213 (light blue, yellow, PDB: 6V05). (C) BamA crosslinks to LptD-ΔD330 at the lateral gate and luminal wall. Positions which only crosslink to LptD4213 are shown in light blue. S439 and Y468 (yellow) crosslink to both LptD4213 and LptD-ΔD330. (D) Positions along β-strand 4 where crosslinking to LptD-ΔD330 or LptD4213 was assessed. (E) The BamA luminal wall crosslinks LptD-ΔD330 less strongly than LptD4213. Weaker crosslinks are denoted with red arrowheads.

Because both LptD4213 and LptD-ΔD330 accumulate near residue E470, we wondered if additional residues in β-strand 4 would also crosslink to these slow-folding substrates. We incorporated *p*Bpa at three other residues (Q466, S472, and T474) whose side chains face the lumen of BamA in β-strand 4 (Figure 2D). We found that all residues (Q466, Y468, S472, and T474) proximal to E470 show UV-dependent crosslinking to LptD4213 and, to a lesser extent, to LptD-ΔD330 (Figures 2E and S2). This is consistent with the observation that LptD-ΔD330 creates a less severe folding defect than LptD4213 as shown by both decreased membrane permeability (Figure 1G), and faster folding as judged by pulse-chase analysis (11). Because LptD mutants accumulate in the lumen of BamA near E470 when they fail to fold, we wanted to know more specifically what regions of the mutant substrates are accumulating in that area of the machine.

### The region of the substrate that accumulates at the luminal wall of BamA is adjacent to the deleted region of the mutant LptD variants

In vivo photocrosslinking does not provide information about what region of the mutant substrate interacts with the luminal surface of BamA. Therefore, we decided to pursue an alternative approach involving the formation of disulfide crosslinks between BamA and mutant LptD substrates (Figure 3A). The longer a region of the substrate to be folded persists in the lumen of BamA, the more likely that a cysteine contained in that region forms a disulfide bond with a cysteine on the BamA luminal wall. We constructed two plasmids each with a cysteine incorporated in either β-strand 4 (Y468, near E470), or β-strand 6 (D512) of BamA (Figure 3B). Additionally, we constructed a series of plasmids encoding LptD4213 with a cysteine incorporated into one of the β-strands spanning from β-strand 2 to β-strand 8. This region of LptD4213 contains the strands which are connected to the deleted loop, as well as the strands which fold immediately after the deleted region in a C-terminal-to-N-terminal folding model (11, 20) (Figures 3C, 3D, S3A, S3B). We grew cells containing both one of the three cysteine-containing BamA and one of the seven cysteine-containing LptD4213. We purified the resulting lysates by pulling down on BamA and probed the presence of high molecular weight crosslinks by immunoblotting.

**Figure 3:**
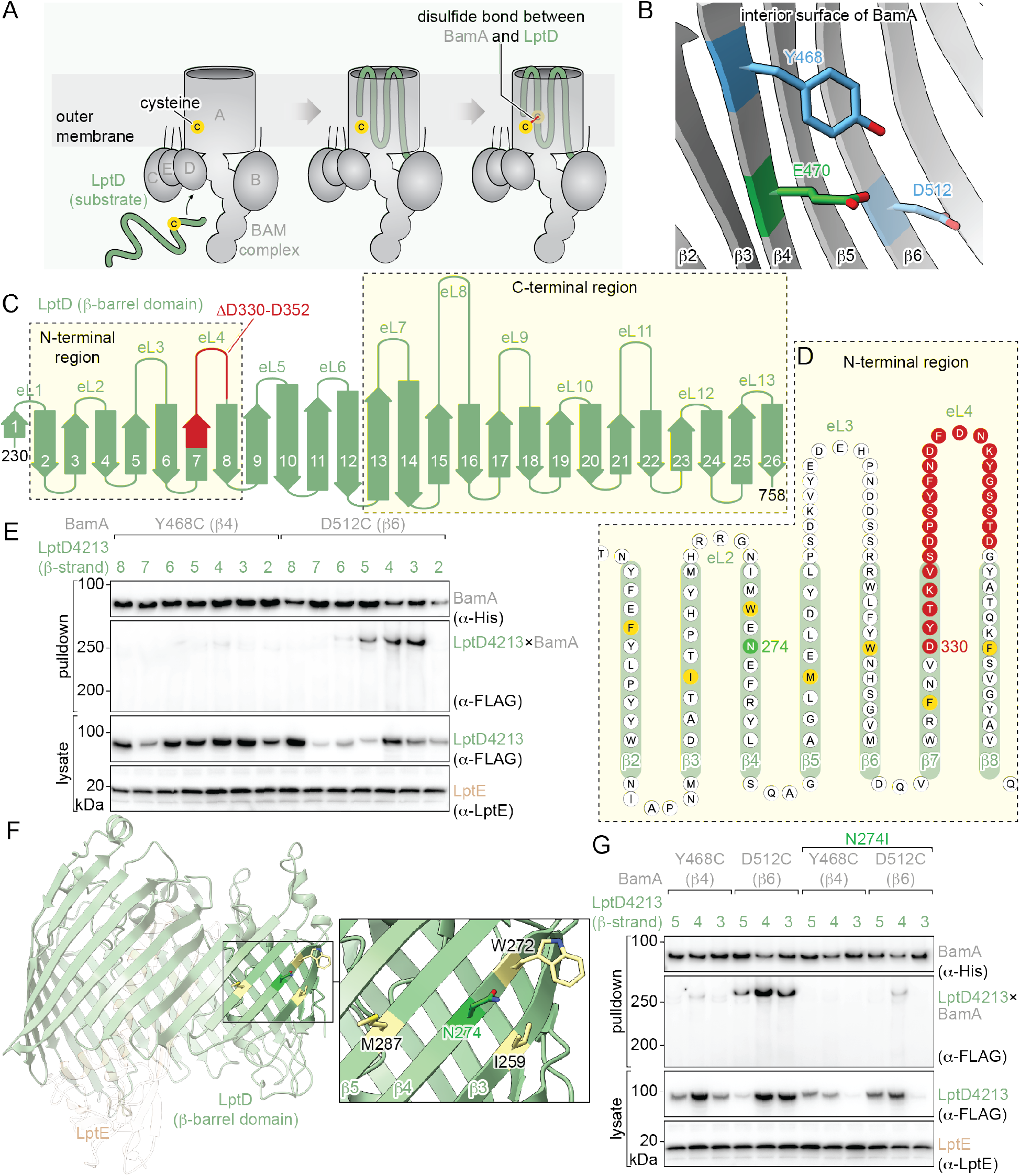
LptD variants accumulate at the BamA luminal wall at a position near the LptD4213 deletion. (A) Disulfide crosslinking assay used to assess the residence time of LptD variant positions within the BamA lumen during folding. (B) Positions Y468 or D512 near E470 in BamA were substituted with cysteine to enable disulfide crosslinking. (C) Diagram of LptD barrel showing the N-terminal region and C-terminal region tested for crosslinking. (D) N-terminal region of LptD showing position of the LptD4213 deletion (red) and N274 (green) as well as positions in each strand tested for disulfide crosslinking to the BamA lumen (yellow). (E) BamA-D512C crosslinks LptD4213 β-strands 5-3 more strongly than BamA Y468C. (F) N274 faces the membrane on strand 4 of LptD, proximal to the residues used for testing disulfide crosslinking. (G) The N274I mutation nearly eliminates the crosslinks between the LptD4213 N-terminal region and BamA D512C.

We found that the cysteine substitution in β-strand 6 of the machine (i.e., BamA-D512C) produced stronger disulfide crosslinks to substrates than that in β-strand 4 (BamA-Y468C) (Figure 3B and 3E). Cysteines in β-strands 5, 4, and 3 of LptD4213 crosslinked strongly to the cysteines placed in the luminal wall of BamA at position D512C (Figure 3E). The observation of crosslinks at these positions, but not at β-strands 8, 7, or 6, which are closer to the deletion ΔD330–D352 in LptD4213, indicates that the cysteine at D512C is reporting on lumen accumulation of backed-up substrate awaiting to be folded. It had not escaped our attention that the cysteines we incorporated in β-strands 5–3 surround residue N274 (Figure 3F). We have previously shown that the mutation N274I is an intragenic suppressor of LptD4213 (35) (Figure S4). If our hypothesis is correct that deletion of the 23 amino acids in the region surrounding β-strand 7 of LptD4213 causes unfolded N-terminal β-strands 5–3 of the substrate to accumulate in the lumen of BamA, incorporation of the intragenic suppressor N274I might decrease the intensity of disulfide crosslinking in that region. We found that incorporation of N274I in LptD4213 nearly eliminated disulfide crosslinking for both reporter positions in the lumen of BamA (Y468 and D512) (Figure 3G). Since mutating this asparagine to an isoleucine alleviates outer membrane defects related to folding of LptD4213 and dramatically diminishes formation of these disulfide crosslinks, we conclude that the strong crosslinks between β-strands 5–3 and the luminal surface of BamA indeed reflect the longer residence time of this region of LptD4213 in the lumen during folding. In essence, LptD4213 becomes trapped in a deep energetic well at this point in the folding process, and the reason that N274I alleviates both folding defects and crosslinking strength is that it destabilizes binding of LptD4213 to the luminal wall of BamA at that same point.

### The folding pathway of wild-type LptD contains two stable intermediates

Since we suggested that LptD-ΔD330 exhibits the same fundamental folding defect as LptD4213 (Figure 1), we wondered whether this region (β-strands 5–3) of other LptD variants also accumulates within BamA during folding. We repeated these experiments incorporating cysteines in β-strands 8–2 of LptD-ΔD330 and wild-type LptD. As observed for LptD4213, LptD-ΔD330 also shows accumulation in the same region, localized to β-strand 4 and β-strand 3 (Figure 4A). Remarkably, even the native substrate shows crosslinking to BamA at this region. The fact that crosslinking at the BamA luminal wall occurs for all three substrates implies that there is an intrinsic barrier at this intermediate step even in the normal folding process. Furthermore, this observation suggests that LptD-ΔD330 and LptD4213 slow the folding by exacerbating this natural energy well. This exciting observation led us to ask whether there are other slow-folding regions within wild-type LptD.

**Figure 4:**
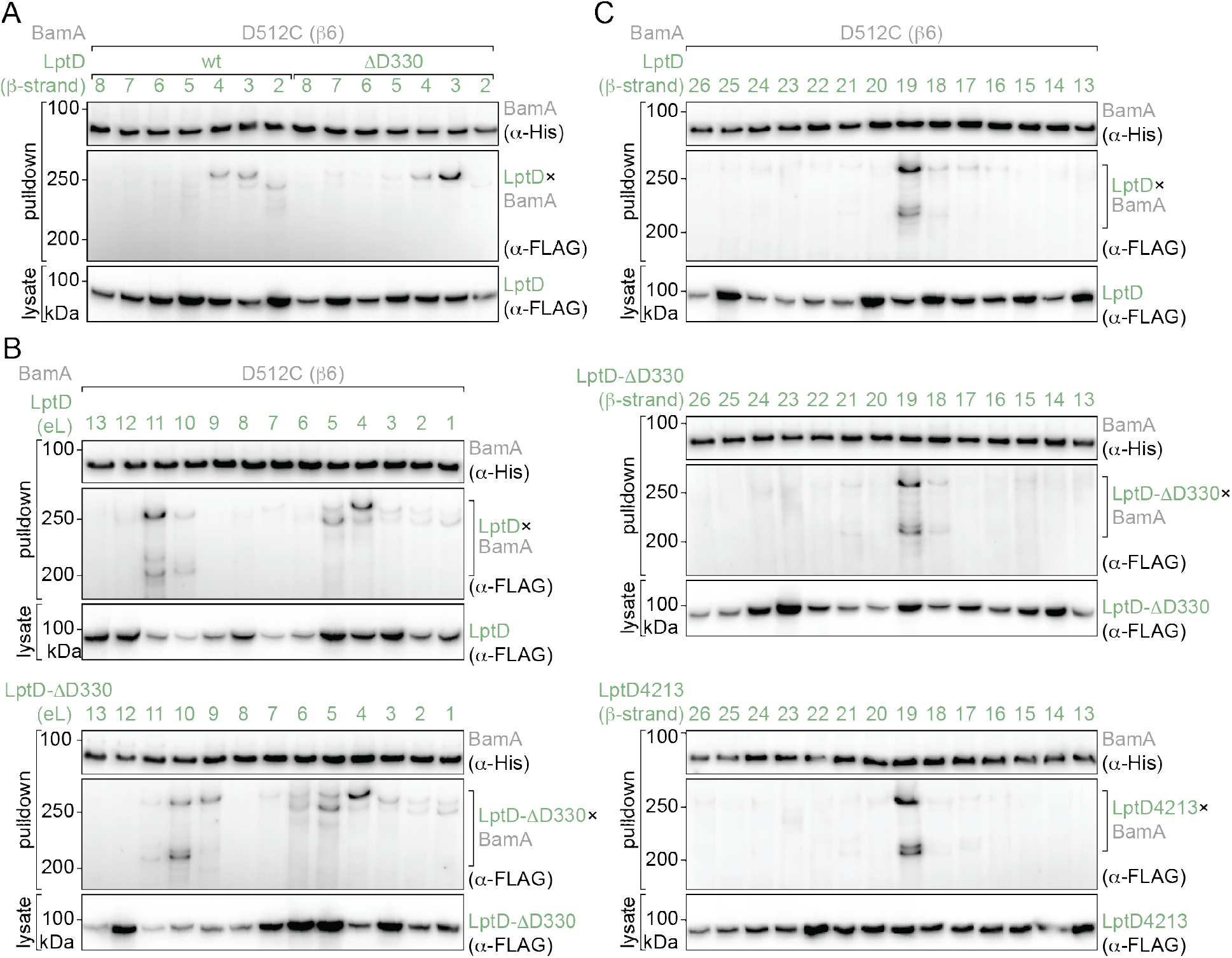
The folding pathway of wild-type LptD contains an N-terminal intermediate and a C-terminal intermediate. (A) β-strands 4–3 in the N-terminal region of both wild-type LptD and LptD-ΔD330 have a long residence in the lumen of BamA. (B) A set of C-terminal loops and a set of N-terminal loops of both wild-type LptD and LptD-ΔD330 have long residence time in the BamA lumen. (C) Disulfide crosslinking between the C-terminal 13 β-strands of wild-type LptD, LptD-ΔD330, or LptD4213 and BamA-D512C. Each variant shows a crosslink within strand 19 of LptD to the BamA luminal wall during folding.

We carried out a comprehensive investigation of the propensity of any of the β-strands and extracellular loops of LptD to reside in the lumen of BamA (Figure 3C, *C-terminal region*, and Figures S3A–C). We found a broad N-terminal region that exhibits crosslinks to BamA-D512C. This region surrounds the extracellular loop that is deleted in LptD4213; it encompasses extracellular loops 5 and 4 (the loop which we have previously established is critical for release of LptD from BamA (11, 33)), as well as β-strands 4 and 3 (Figures 4A and 4B). We found a separate earlier region of wild-type LptD, encompassing β-strands 19 and extracellular loops 11 and 10, that crosslinks strongly to position D512C of BamA (Figures 4B and 4C). Remarkably, this region of LptD folds long before the N-terminal region which we have discovered contains a natural barrier to folding. The two slow-folding LptD substrates (LptD-ΔD330 and LptD4213) show the same crosslinking intensity as observed for folding of the wild-type substrate (Figures 4B and 4C). This observation is consistent with our findings that the mutations affect the energy landscape at the N-terminal intermediate rather than the C-terminal intermediate. Thus, it appears that there are at least two observable folding intermediates for LptD: one N-terminal intermediate and one C-terminal intermediate.

### β-barrel substrates have common folding intermediates

We wondered whether other β-barrel substrates exhibit long-lived intermediates like the ones we observed for LptD. We carried out an experiment analogous to the one described above to examine the residence time of a BamA substrate (BamA^S^) on the BamA machine (BamA^M^) (Figure 5A). Our BamA^S^ lacks a portion of its N-terminal soluble periplasmic polypeptide transport-associated (POTRA) domain (ΔG172–R421) to prevent lipoprotein binding upon substrate folding. However, unlike a previously studied slow-folding BamA^S^ that has a deletion of residues in extracellular loop 1 (I430–S439) (20), the substrate we used here contains no deletions in the β-barrel domain that would alter the folding pathway of BamA^S^.

**Figure 5:**
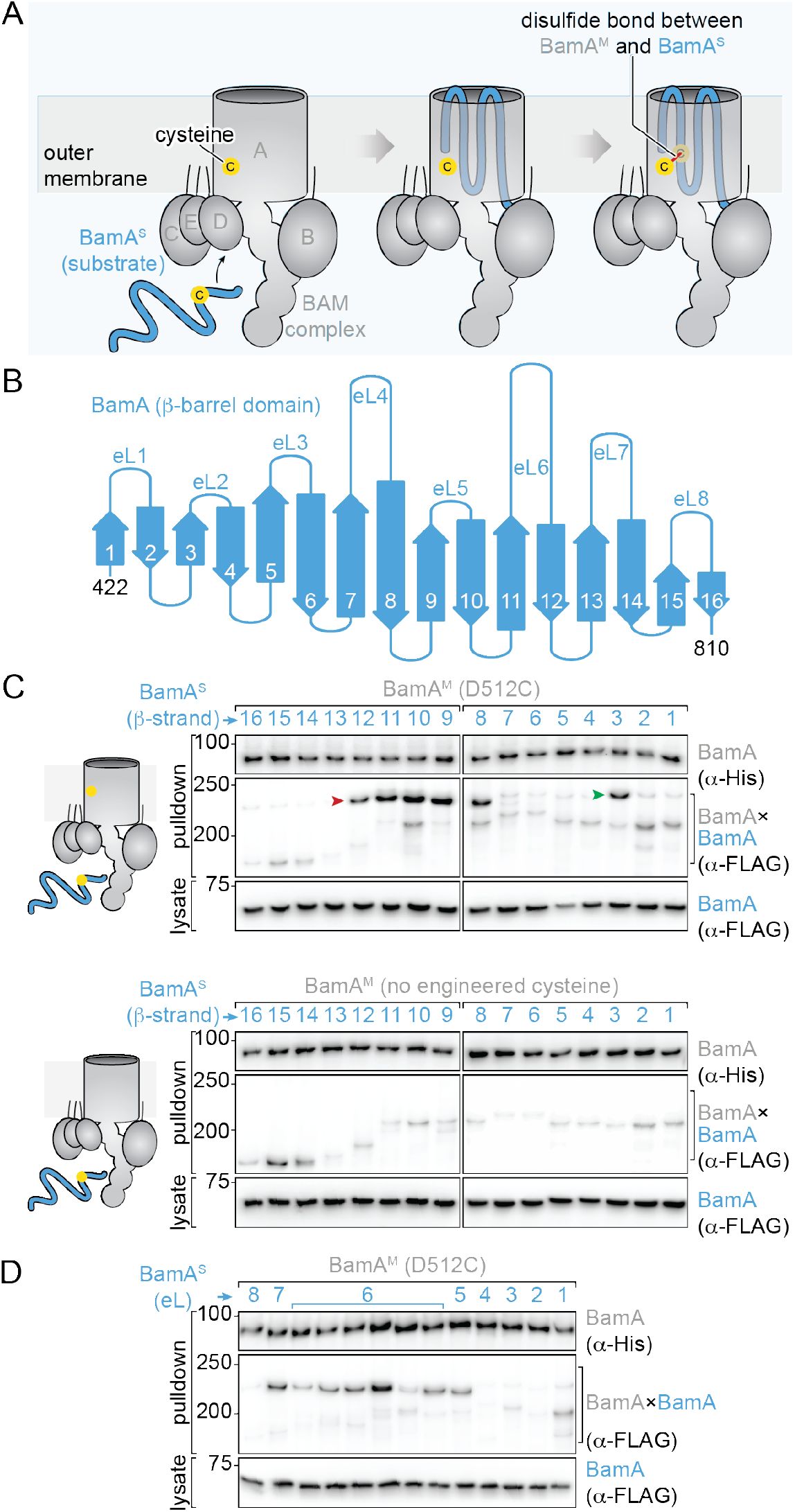
BamA^S^ exhibits folding intermediates at the same points in folding as LptD. (A) Disulfide crosslinking assay used to assess residence time of BamA^S^ strands or loops within the lumen of BamA^M^ during folding. (B) Diagram of the 16-stranded BamA^S^ barrel. (C) Disulfide crosslinking between each β-strand 16–1 of BamA^S^ and the BamA^M^ luminal wall with or without D512C in BamA^M^. Two sets of BamA^S^ strands crosslink to the BamA^M^ luminal wall: β-strands 12–8 and β-strand 3. These crosslinks depend on the presence of BamA^M^-D512C. (D) Disulfide crosslinking between positions within each extracellular loop of BamA^S^ and the BamA^M^ luminal wall. Crosslinking positions fall within the C-terminal loops 7–5 of BamA^S^.

We introduced cysteines individually in each of the 16 β-strands and the eight extracellular loops of BamA^S^ and asked whether specific parts of the substrate display a long residence time proximal to position D512C of the BamA^M^ (Figures 5A and S5). Although we detected bands corresponding to the formation of disulfide bonds for all 16 β-strands of the BamA^S^, two regions display strong disulfide crosslinks (Figure 5C, *left panel*). One region comprises β-strands 12–8 (Figure 5C, *left panel, red arrow*) and the other region, β-strand 3 (Figure 5C, *left panel, green arrow*). As a control, we performed a similar experiment using a BamA^M^ that does not contain the reporter cysteine at position D512 (Figure 5C, *right panel*) and found that all the weak crosslinks we had observed persist. However, the two regions of BamA^S^ that strongly crosslink to the BamA^M^ containing a cysteine at position D512C completely disappear. This result indicates that the strong crosslinks we observed at these two regions of BamA^S^ are dependent on the engineered cysteine at position D512. Furthermore, we also observed the formation of strong crosslinks between extracellular loops 7–5 of BamA^S^ and BamA^M^-D512C (Figure 5D). Taken together, these results suggest that, like LptD, BamA^S^ has two observable intermediates in its natural folding process: one near the C-terminus of the substrate and one near the N-terminus of the substrate. Of note, the C-terminal intermediate appears approximately 7 strands from the C-terminus in both BamA (β-strand 9) and LptD (β-strand 19). Similarly, the N-terminal intermediate appears approximately 3-4 strands from the N-terminus in both substrates.

### BamA-E470 variants increase substrate binding to the BAM machine

We have previously suggested that the interior surface of BamA catalyzes folding of β-barrel substrates (11, 20, 33). Furthermore, we found that BamA-E470G is a variant that restores the rate of folding of a slow-folding mutant of LptD. We also know that the intragenic suppressor, N274I, of LptD4213 eliminates the intense crosslinks formed by this slow-folding substrate at β-strands 5, 4, and 3. Therefore, we wondered whether mutations at E470 of BamA similarly decrease the residence time of LptD-ΔD330 in the lumen at the N-terminal slow step of folding. We examined the effect of the extragenic suppressor E470G on the disulfide crosslinks between BamA-D512C and LptD-ΔD330 that we had previously observed. Surprisingly, the E470G mutation in BamA did not decrease the crosslink intensity at strands 4 or 3. If anything, the intensity was slightly increased (Figure 6A). BamA-E470G increases the rate of folding of LptD-ΔD330 from about 50 minutes to about 25 minutes (11), but does not apparently alleviate the N-terminal energy well which is exacerbated by the deletion of residues D330. This observation requires a different explanation for how altering E470 increases folding rate.

**Figure 6:**
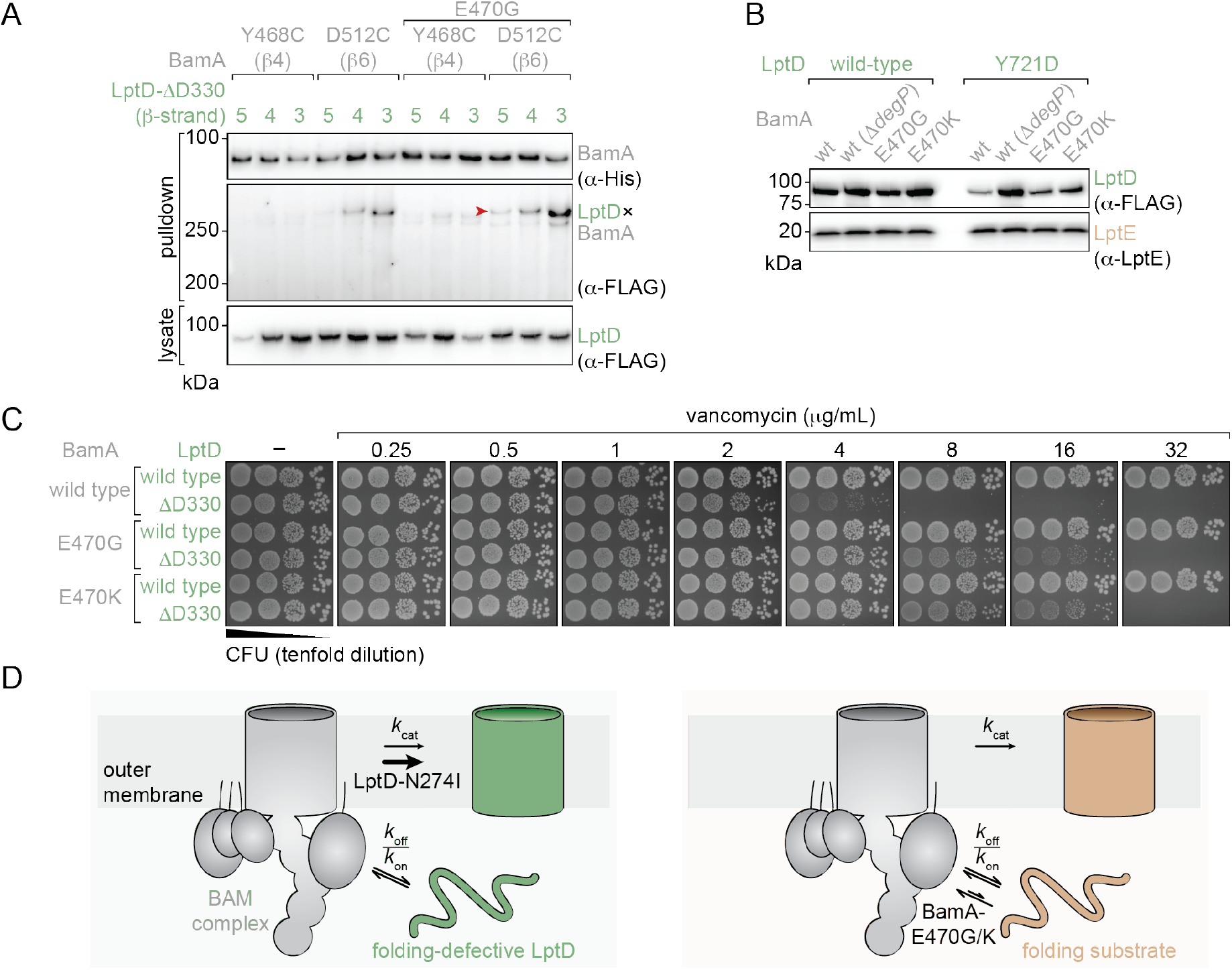
Mechanism of substrate rescue by BamA-E470G/K variants. (A) Disulfide crosslinking between the N-terminal strands of LptD-ΔD330 and the luminal wall of BamA with or without E470G. BamA-E470G does not decrease LptD-ΔD330 binding during the late folding intermediate. (B) Levels of folded wild-type LptD and LptD-Y721D variants in cells in the presence of chromosomal copies of BamA E470 variants or DegP deletion. E470G and E470K partially rescue the folding of LptD Y721D. (C) Vancomycin sensitivity assay of *E. coli* containing LptD-ΔD330 in cells with a chromosomal copy of wild type BamA, BamA-E470G or BamA-E470K. BamA-E470G and BamA-E470K similarly rescue the membrane permeability defect imparted by LptD-ΔD330 (D) Model for the differential activity of the suppressors LptD-N274I and BamA-E470G/K. LptD-N274I is proposed to decrease the affinity of BAM for defective LptD substrate during the late folding intermediate, thereby increasing the folding rate by increasing *k*_*cat*_ (left panel). Conversely, BamA-E470G/K is proposed to increase OMP flux by increasing the affinity of the BamA for substrate at the start of folding, decreasing *k*_*off*_/*k*_*on*_ (right panel).

Another variant of BamA-E470, BamA-E470K, has previously been identified as a bypass suppressor of the essentiality of BamD (36). We had previously identified BamD as the binding site for the targeting β-signal conserved among β-barrel substrates of the BAM complex (18). The binding of BamD to unfolded substrates and the subsequent loading of substrates onto BamA is likely part of the essential function of BamD. Thus, since BamA-E470G did not appear to decrease the residence time of LptD-ΔD330 in the lumen, and since another variant at that same position on the luminal wall bypasses the essential function of a protein that binds to unfolded substrates, we hypothesized that the E470G suppressor accelerates the rate of folding of LptD-ΔD330 by increasing its binding to BamA.

To test this hypothesis, we utilized a variant of LptD containing the Y721D mutation in the β-signal of LptD. We have previously shown that LptD-Y721D binds BamD, but does not readily advance to BamA and, therefore, is removed from BAM by Skp and degraded by the periplasmic protease DegP rather than being effectively folded into the membrane (19, 37). We concluded that a competition exists: the slower the loading of a substrate onto BamA and its integration into the outer membrane, the more exposure the substrate has to degradation in the periplasm by the Skp/DegP pathway. We engineered *E. coli* strains that contained either a chromosomal copy of the gene encoding BamA-E470G or BamA-E470K and assessed the levels of either wild-type LptD or LptD-Y721D in the presence of each of these mutations. We found that both BamA-E470G and BamA-E470K partially rescued the substrate from periplasmic degradation by DegP (Figure 6B). We conclude that these mutations at the luminal wall of BamA increase LptD-Y721D binding to BamA which increases the likelihood that the substrate will fold rather than be degraded.

Finally, we tested the ability of BamA E470K to rescue membrane permeability defects caused by LptD-ΔD330. We found that BamA-E470G and BamA-E470K identically rescued cells containing LptD-ΔD330 from vancomycin (Figure 6C). These results provide insight into the role that the region surrounding BamA-E470 plays in the folding process. Unlike the intragenic suppressor LptD4213-N274I which alters the folding pathway of LptD4213 to lower the barrier surrounding LptD4213 β-strands 5–3, the extragenic suppressors BamA-E470G and BamA-E470K rescue folding of LptD-ΔD330 by increasing substrate binding to the luminal wall of BamA. In this way, they affect the rate of folding by altering the equilibrium of substrate binding to BamA and thus the throughput of substrate, rather than changing the depth of the well at the N-terminal intermediate.

## Discussion

In this work we identified two intermediates in the folding of two wild-type β-barrels, LptD and BamA, as they fold on the BAM complex. In LptD, the first of these intermediates exists during assembly of β-strand 19 and extracellular loops 11 and 10. In BamA, the first of these intermediates comprises the assembly of β-strands 12 through 8. Because LptD has 26 β-strands, BamA has 16 β-strands, and folding of substrates starts at the C-terminus, the first intermediate of folding for each of these substrates appears at the same stage: about seven strands into the folding process. Similarly, the second intermediate exists during assembly of LptD β-strands 4–2, and BamA β-strand 3. The barrier responsible for this second intermediate in folding is therefore also shared because it occurs around three strands from completion of the folding process for both LptD and BamA. This observation is consistent with BAM catalyzing the folding of a variety of β-barrel substrates which vary in size from 8 to 36 β-strands (38, 39). Further characterization of these shared intermediates will allow us to understand how the BAM complex addresses general membrane protein folding challenges.

We also characterized two slow-folding variants of LptD, LptD4213 and LptD-ΔD330, which share a common folding defect. We have characterized the behavior of suppressors for each of these slow-folding variants. Several years ago, the intragenic suppressor of LptD4213, LptD4213-N274I (previously known as Imp61), was shown to substantially alleviate membrane permeability defects and allowed a much larger percentage of properly folded substate to form (33, 35). Here, we show that this suppressor restores the second, exacerbated barrier in LptD4213 folding to wild-type levels.

BamA-E470G does not alter the late barrier observed for LptD-ΔD330 like N274I does for LptD4213. the suppressors LptD-N274I and BamA-E470G must restore the rate of folding in different ways. We wanted to understand how BamA-E470G could restore the folding rate of a defective substrate without apparently affecting either of the barriers that we have shown to exist. A first clue came from the fact that BamA-E470K has been previously characterized as a bypass suppressor of the essentiality of BamD, allowing cells lacking BamD to be viable (36). It has been argued that this suppressor does not change the catalytic activity of the BAM complex, but rather bypasses a checkpoint at which BamD would normally give approval to fold. Before substrates begin to fold on BamA, BamD binds to their β-signal: a conserved motif of β-barrel substrates near the C-terminus. It has been shown that β-signal mutants such as LptD-Y721D have a faster off-rate because they do not transfer to BamA as well as the native substrate (19, 37).

A second clue came from previous identification of E470G as a mutation which restores folding activity to a defective BamA chimera made from the *E. coli* β-barrel domain and four of the five *P. aeruginosa* POTRA domains (40). Since the POTRA domains coordinate the arrangement of the four lipoproteins, including BamD, the folding defect of this chimera likely stems from inability of substrates to effectively get on BamA via BamD and start folding. Together, these clues suggest that BamA-E470G and BamA-E470K work similarly by providing a second alternative pathway for substrates to bind to BamA and start folding without binding BamD.

We propose the following model to explain how LptD-N274I and BamA-E470K/G differently increase folding rates of defective substrates (Figure 6D). Fundamentally, there are two ways to increase the folding rate of LptD4213 and LptD-ΔD330. Since both mutants increase the barrier associated with the N-terminal intermediate step of folding, a suppressor could lower that barrier by decreasing the binding of substrate to the machine during that intermediate step. This strategy appears to be how LptD-N274I restores the folding rate of LptD4213. LptD-N274I thus alleviates membrane permeability defects by freeing up available BAM machines which would otherwise be jammed with stalled substrate. However, the BamA-E470 variants do not lower the N-terminal barrier observed during folding. They restore wild-type rates of folding by providing an alternate pathway for substrate binding to BamA in the initial stage of folding, thereby decreasing *k*_off_ from BamA. These variants likely alleviate the outer membrane permeability defects of LptD-ΔD330 by generally increasing the flux of OMPs, including other BamA molecules, into the outer membrane.

The similarity of BamA-E470G and BamA-E470K implies that in addition to being a selectivity filter, BamD accelerates folding by directly facilitating substrate engagement with BamA. We therefore propose that an editing function exists for the three proteins BamD, Skp, and DegP. Together, these proteins create the requirement for a β-signal: a proper substrate with a β-signal will bind to BamD, transfer to BamA, and begin to fold, hence avoiding the Skp/DegP-mediated degradation pathway. A substrate without a proper β-signal will not transfer to BamA, get removed from BamD by Skp, and be exposed to periplasmic degradation by DegP.

Finally, a hallmark of the LptD4213 mutant is the presence of severe cell permeability defects. We have shown here that the slow-folding substrate LptD4213, and to a lesser extent LptD-ΔD330, gets trapped in the late folding intermediate state on the BAM complex. Because the folding of wild-type LptD also requires passing through this late intermediate, further understanding of the late folding intermediate could enable the design of drugs that permeabilize the Gram-negative outer membrane.

## Materials and Methods

### Bacterial strains and growth conditions

We cultivated all bacterial strains aerobically at 37°C in lysogeny broth [LB, 1% (w/v) tryptone, 0.5% (w/v) yeast extract, and 1% (w/v) NaCl] or in LB containing 1.5% (w/v) agar. When necessary, we added carbenicillin (50 mg/mL), chloramphenicol (25 mg/mL), kanamycin (30 mg/mL), spectinomycin (50 mg/mL), vancomycin (various concentrations, as described in the manuscript), arabinose (0.05% or 0.2%), and glucose (0.05% or 0.2%) to the growth media. All bacterial strains and plasmids used in this study are listed in Supplementary Tables 1 and 2, respectively.

### Genetic methods

We used standard procedures for genomic and plasmid DNA purification, amplification, mutagenesis, cloning, and transformation. Genomic DNA was isolated with the Wizard Genomic DNA Purification Kit (Promega). KOD Hot Start DNA polymerase (MilliporeSigma) or Q5 High-Fidelity DNA Polymerase (New England Biolabs) were used to generate entire plasmids or DNA fragments for cloning and mutagenesis, and OneTaq DNA polymerase (NEB) was used for general analytical PCR. We used a modified chemical method (41) to prepare and transform competent *Escherichia coli* cells. Plasmids were isolated using the alkaline lysis method, followed by silica-based purification, and verified by sequencing. We used allelic exchange (42) to introduce chromosomal modifications (nucleotide replacements and deletions) and verified all changes by PCR analysis, DNA sequencing, and by testing susceptibility to antibiotics. All oligonucleotide primers used in this study are listed in Supplementary Table 3.

### SDS-PAGE and immunoblotting analysis

Purified proteins and protein mixtures from cell lysates were analyzed on 4–8% or 4–20% polyacrylamide gradient Tris-HCl gels at 150 V for 1 h in Tris-glycine-SDS running buffer [25 mM Tris-HCl (pH 8.0), 192 mM glycine, 0.1% SDS (w/v)]. The 2× SDS sample buffer contained 125 mM Tris (pH 6.8), 4% (w/v) SDS, 30% (v/v) glycerol, and 0.005% bromophenol blue. Following electrophoresis, gels were either stained with a solution of Coomassie brilliant blue [0.1% (w/v) Coomassie Blue R-250, 10% (v/v) acetic acid, and 50% (v/v) methanol] or used for immunoblotting analysis. For immunoblot analysis, we transferred proteins onto Immun-Blot polyvinylidene difluoride (PVDF) membranes (BioRad) using a Power Blotter semi-dry transfer system (Invitrogen) and incubated the membranes in 1× Casein Blocking Buffer (Sigma-Aldrich) diluted in 1× TBST buffer [20 mM Tris-HCl (pH 8.0), 137 mM NaCl, 1% (v/v) Tween-20)] at 25°C for 1 hour. Primary antibodies were diluted in 1× TBST and the membranes were incubated in this solution at 4°C overnight. We washed the membranes three times in 10 mL of 1× TBST at 25°C for 10 min. Secondary antibodies were diluted in 1× TBST and the membranes were incubated in this solution at 25°C for 1 h. We washed the membranes three times in 10 mL of 1× TBST at 25°C for 10 min and developed them for chemiluminescence using the Amersham ECL Prime Western Blotting Detection Reagent (Cytiva). Immunoblots were imaged using an Azure Biosystems C400 imager. Antibody dilutions were as follows: anti-LptD at 1:10,000, anti-LptE at 1:10,000, Direct-Blot HRP anti-6×His Epitope Tag antibody (BioLegend) at 1:10,000, Direct-BlotHRP anti-DYKDDDDK Tag (BioLegend) at 1:10,000, and HRP-conjugated goat anti-rabbit IgG (Thermo Fisher Scientific) at 1:10,000.

### Antibiotic sensitivity assay

To measured bacterial sensitivity to vancomycin, we grew cell cultures with agitation (200 rpm) at 25°C overnight. Saturated overnight cultures were diluted (1:100) into 2 mL of fresh LB and incubated at 37°C with agitation (200 rpm) until reaching an optical density at 600 nm (OD_600_) of 0.3 ± 0.02 (about 1×10^8^ colony-forming unit per mL, CFU/mL). Were serially diluted the calibrated cultures in 1× PBS (137 mM NaCl, 2.7 mM KCl, 10 mM Na_2_HPO_4_, 1.8 mM KH_2_PO_4_, pH 7.4) and plated 10-μL aliquots of each dilution onto LB agar and LB agar containing increasing concentration of vancomycin. The plates were incubated at 37°C for 16–20 h. For experiments involving the conditional expression of LptD, we added arabinose or glucose to the culture media to induce or repress, respectively, the expression of LptD.

### Site-specific in vivo photo-crosslinking of BamA to substrates

The method for genetic incorporation of the photoreactive, unnatural amino acid *p*-benzoyl-L-phenylalanine (*p*Bpa) at an amber codon engineered into specific positions of the gene encoding BamA was adapted from previous work (11, 43). Cells harboring the plasmids pSup-BpaRS-6TRN and pZS21-6×His-*bamA* containing individual amber mutation at the indicated positions were grown at 37°C overnight, diluted (1:100) into 100 mL of fresh LB supplemented with 0.8 mM *p*Bpa, and incubated at 37°C with agitation (200 rpm) until reaching an OD_600_ of about 0.8 ± 0.02. Each culture was divided into two aliquots of 45 mL and one of the two aliquots was directly irradiated with UV light (UVP Blak-Ray B-100AP high-intensity UV inspection lamp) at 365 nm at 25°C for 10 min. We harvested the cells by centrifugation (5,000 *g*, 10 min, 4°C), froze the pellets in liquid nitrogen, and stored them at -80°C.

### Site-specific in vivo disulfide crosslinking

Cells harboring the plasmids pZS21-6×His-*bamA* and pTrc99A-*lptD*-3×FLAG (or one of the variants, Supplementary Table 2) containing engineered cysteines into specific positions of the genes encoding BamA and LptD were grown at 37°C overnight, diluted (1:100) into 25 mL of fresh LB and incubated at 37°C with agitation (200 rpm) until reaching an OD_600_ of about 0.6 ± 0.02. At this step, 1-mL aliquots of the cell cultures were collected to monitor the expression levels of the BAM substrates LptD and BamA^S^. We harvested the cells in the aliquots by centrifugation (21,130 *g*, 2 min, 4°C), resuspended the pellets in 100 μL of BugBuster (MilliporeSigma), incubated the suspension on ice for 10 min, added 100 μL of 2× SDS sample buffer containing 2% (v/v) β-mercaptoethanol. We harvested the remainder of the cells in the original culture by centrifugation (5,000 *g*, 10 min, 4°C), froze the pellets in liquid nitrogen, and stored them at -80°C.

### Protein purification

The method for purification of crosslinked proteins was adapted from previous work (11). Samples were kept at 4°C throughout the entire procedure. We resuspended the pellets in 5 mL of 1× TBS [20 mM Tris-HCl (pH 8.0), 150 mM NaCl] containing 1% (w/v) Anzergent 3-14 (Anatrace), 20 mM imidazole (pH 8.0), 100 μg/mL lysozyme (Sigma-Aldrich), 1mM phenylmethylsulfonyl fluoride (PMSF, Sigma-Aldrich), and 50 μg/mL DNase I (Sigma-Aldrich) and lysed the cells by sonication (25% amplitude, 10 sec on and 10 sec off for a total cycle of 3 min). The cell lysate was incubated with Ni-nitrilotriacetic acid (NTA) agarose on a rocking platform at 4°C for 1 h. We discarded the unbound protein fraction and washed the resin three times with 25 column volumes of 1× TBS containing 0.02% Anzergent 3-14 and 20 mM imidazole (pH 8.0). The bound protein fraction was eluted with 5 column volumes of 1× TBS containing 0.02% Anzergent 3-14 and 200 mM imidazole (pH 8.0). We added TCA (10% of the final volume) to the eluates and incubated the mixture on ice for 20 min. We pelleted the precipitated proteins by centrifugation (21,130 *g*, 10 min, 4°C) and resuspended them in 40 μL (samples from site-specific in vivo photo-crosslinking) or 26 μL (samples from in vivo disulfide crosslinking) of a 1:1 mixture of 2× SDS sample buffer:1 M Tris-HCl (pH 8.0). For samples from site-specific in vivo photo-crosslinking, we added 2% (v/v) β-mercaptoethanol to the 2× SDS sample buffer. After incubating the samples at 95°C for 10 min, we analyzed them by SDS-PAGE and immunoblotting as described above.

## Acknowledgments

We thank N. Ruiz her feedback on the manuscript. This research was supported by the Howard Hughes Medical Institute (Hanna H. Gray Postdoctoral Fellowship to T.M.A.S.) and National Institutes of Health (through award 5R01AI081059-13 to D.E.K.).

## Notes

### Competing Interest Statement

The authors have declared no competing interest.

