## Supplementary Material for "Native β-barrel substrates pass through two shared intermediates during folding on the BAM complex"

**Affiliation:**
*Department of Chemistry and Chemical Biology, Harvard University, Cambridge, MA 02138, USA*

### Supplementary Figures

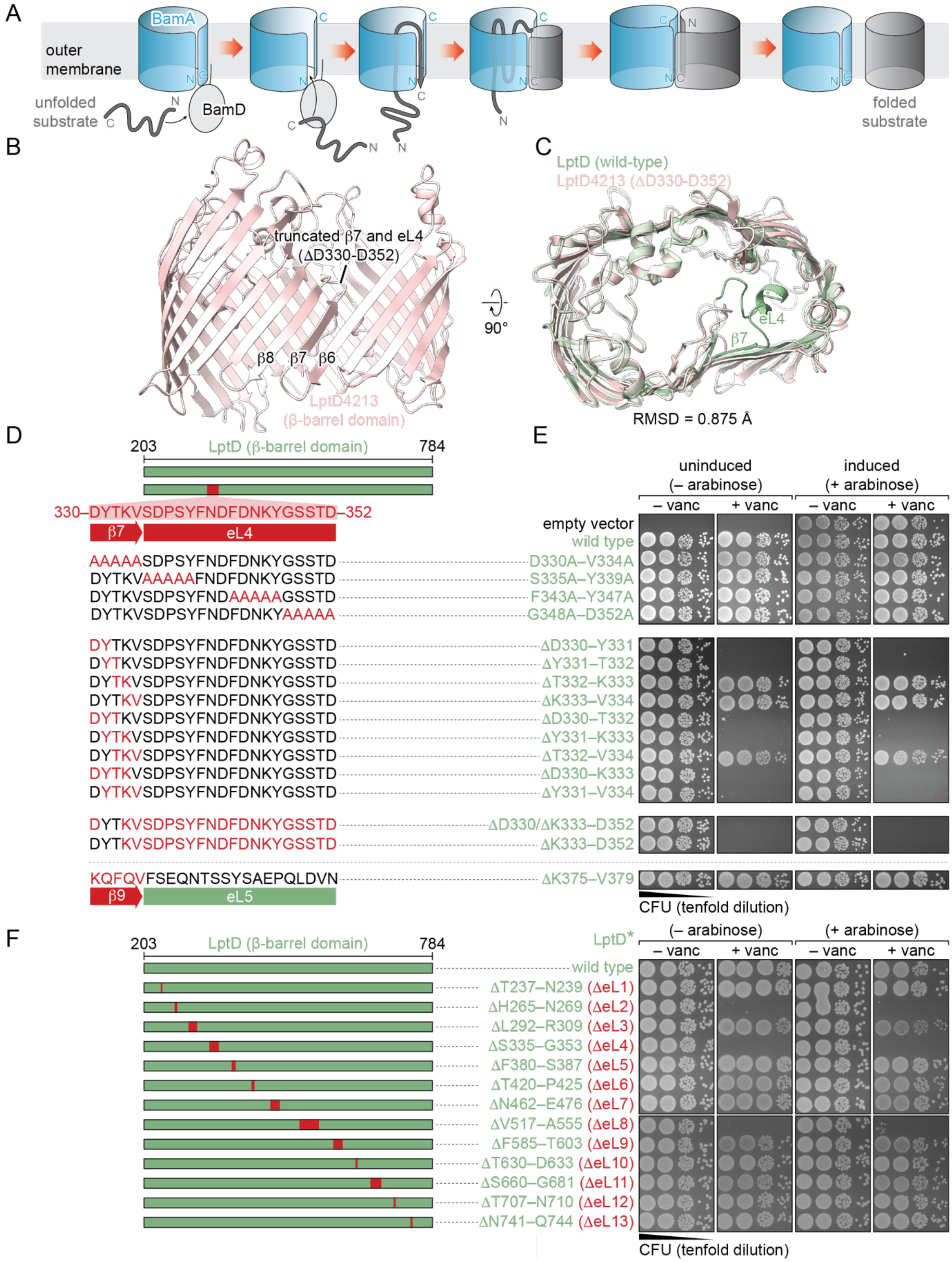

**Figure S1: Register change alone does not recapitulate the folding defects present in LptD4213.** (A) Model of Bam-mediated folding. The outer membrane protein (OMP) substrate β-signal binds to BamD, then BamA. Folding occurs inside the BamA lumen, and folded substrate exits the BamA barrel through the lateral gate. Release occurs once substrate has fully exited the BamA lumen. (B) Model of LptD4213 β-barrel domain produced by Alphafold (1). (C) Overlay of Alphafold-predicted model of LptD4213 β-barrel domain and LptD β-barrel domain (PDB: 4RHB). The overall architecture of the LptD4213 β-barrel domain produced by Alphafold is similar to the experimental structure of the LptD β-barrel. (D) Additional deletions and alterations to the characteristic LptD4213 deletion which were tested in their ability to cause vancomycin permeability defects using a vancomycin permeability assay. (E) Vancomycin sensitivity assay of E. coli containing plasmid A, encoding LptD with deletions shown in panel D, and plasmid B on plates with and without arabinose or vancomycin. Deletion of groups of two, three or four amino acids within the group of LptD D330–V334 can result in permeability defects. This implies that a simple change in register that results from the deletion of D330 is not sufficient to explain the imparted folding defect. (F) Vancomycin sensitivity assay of *E. coli* containing plasmid A, encoding LptD with deletions of extracellular loops 1 through 13, and plasmid B on plates with and without arabinose or vancomycin. Deletion of LptD loops 2, 4, or 8 result in dominant cell permeability defects.

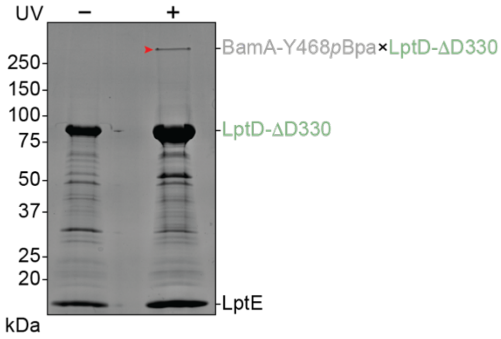

**Figure S2: BamA-Y468*p*Bpa forms a UV-dependent crosslink to LptD-ΔD330.** BamA-Y468*p*Bpa-LptD-ΔD330 crosslinking sample (Figure 2E) was run on an SDS-PAGE gel and imaged using Coomassie staining. There is a resulting UV-dependent high-molecular weight band due to this crosslink (red arrowhead).

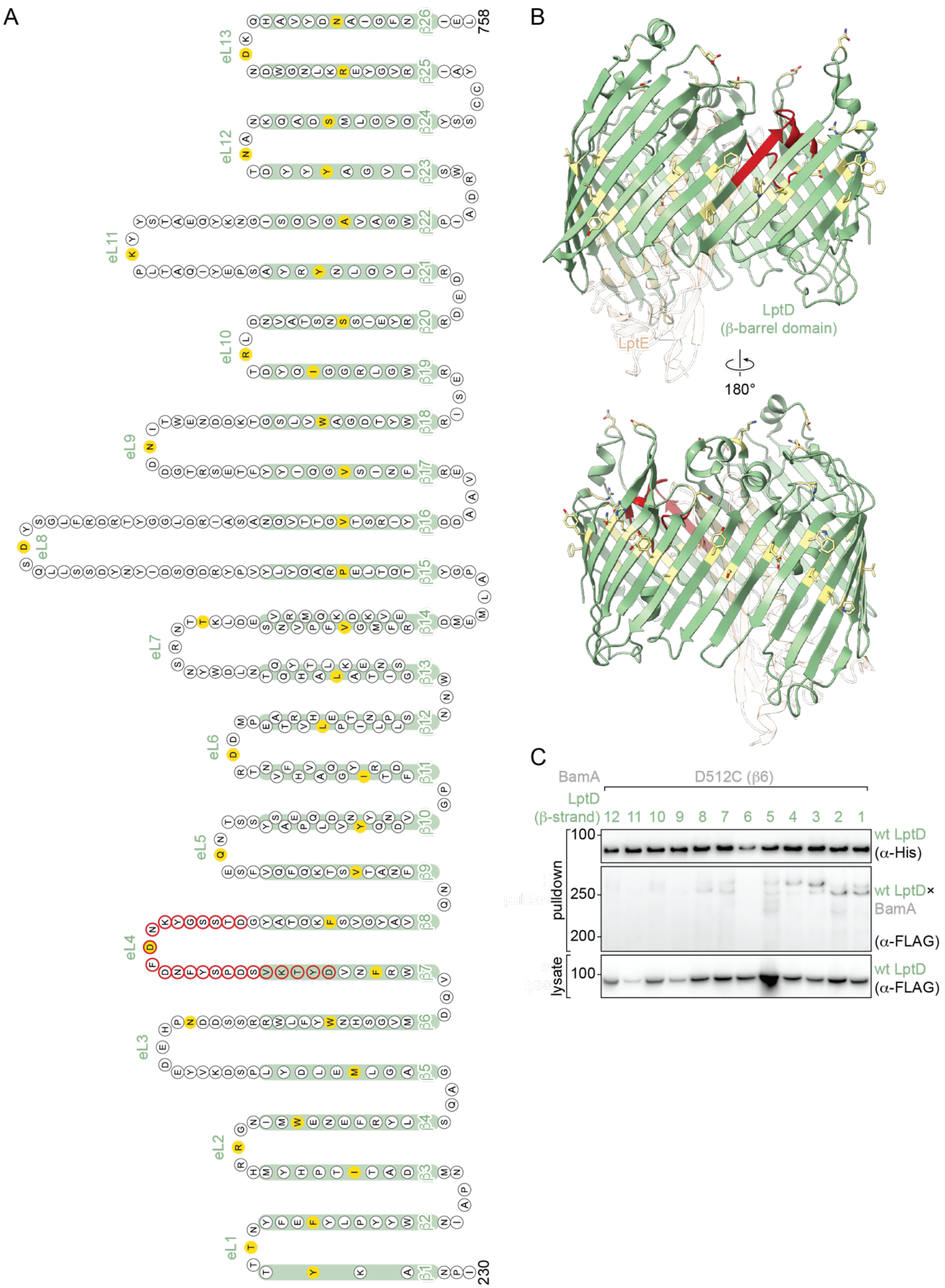

A
B
C
758
230
eL1
eL2
eL3
eL4
eL5
eL6
eL7
eL8
eL9
eL10
eL11
eL12
eL13
LptD
(β-barrel domain)
LptE
180°
BamA
D512C (β6)
LptD
(β-strand)
12 11 10 9 8 7 6 5 4 3 2 1
pulldown
lysate
100
250
200
100
wt LptD
(α-His)
wt LptD×
BamA
(α-FLAG)
wt LptD
(α-FLAG)

**Figure S3: LptD substrate disulfide crosslinking positions.** (A) Diagram showing sequence of β-barrel domain of LptD with the position of the characteristic LptD4213 deletion outlined in red. The positions of cysteine substitutions in each strand and loop used for LptD disulfide crosslinking experiments are shown in yellow. Sequence identities of cysteine substitutions are listed in Table S4. (B) Ribbon model of the LptD β-barrel domain with lipoprotein plug LptE (PDB: 4RHB). The portion of the barrel deleted in LptD4213 is shown in red. Residues chosen for cysteine substitution for disulfide crosslinking experiments are shown as yellow sticks. Residues within β-strands that were chosen for cysteine substitution have outward-facing side chains and are positioned approximately in the middle of each β-strand. Residues within extracellular loops that were chosen for cysteine substitution are positioned approximately in the middle of each loop. C) Disulfide crosslinking between BamA-D512C and wild-type LptD with cysteines substituted in strands 12–1. The C-terminal intermediate encompasses crosslinks between strands 4–2 of LptD and BamA-D512C on the BamA luminal wall.

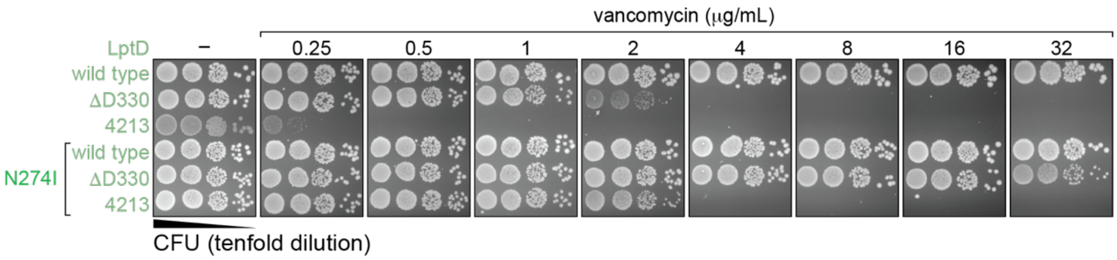

**Figure S4: LptD N274I alleviates folding defects of both LptD-ΔD330 and LptD4213.** Plating assay of *E. coli* containing wild-type LptD, LptD-ΔD330 or LptD421, along with the respective N274I variants, on LB-agar containing varying concentrations of vancomycin. N274I rescued the growth of cells containing LptD-ΔD330 and LptD4213 at high concentrations of vancomycin.

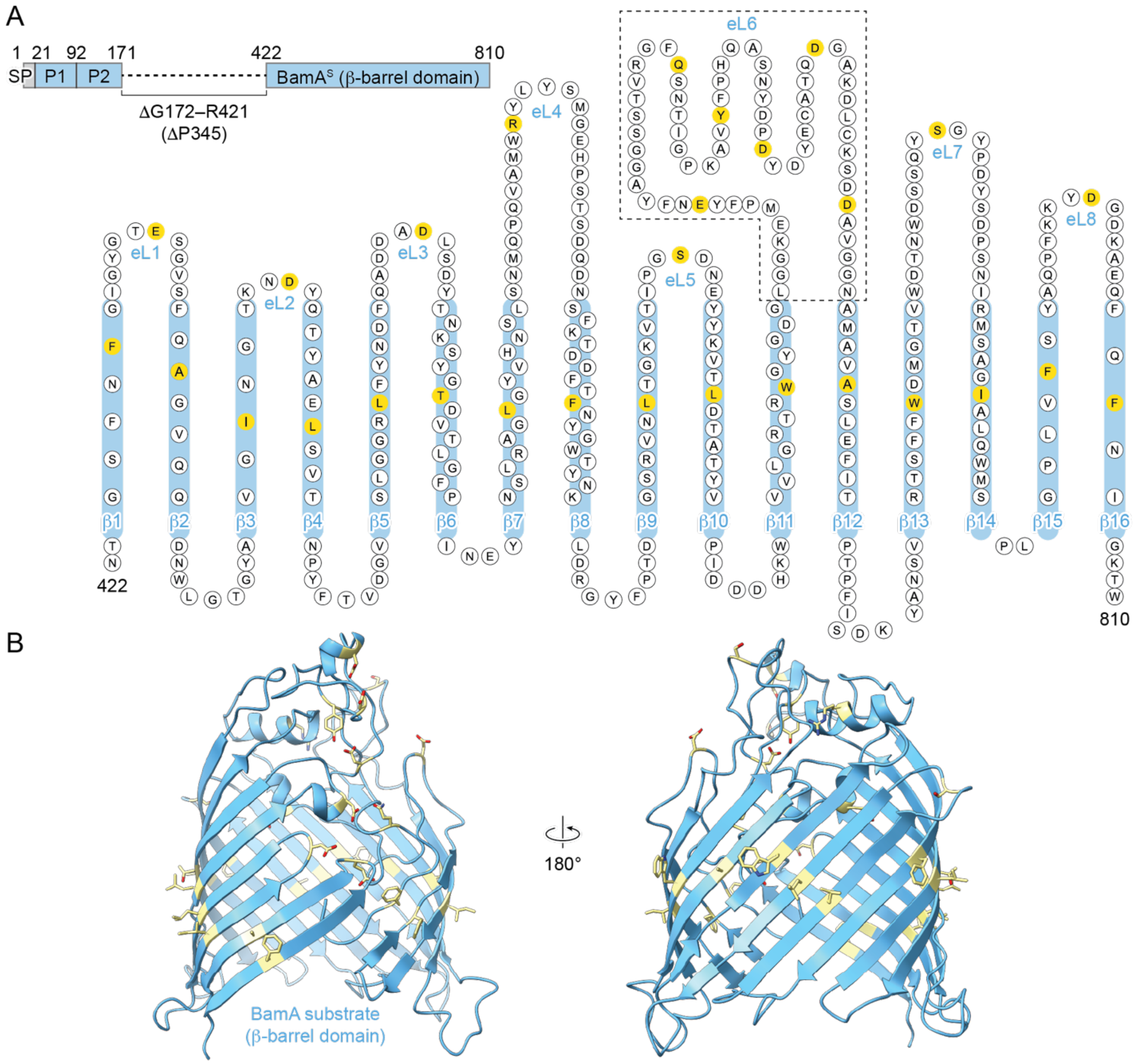

**Figure S5: BamA$^S$ substrate disulfide crosslinking positions.** (A) Diagram showing organization of the β-barrel domain of BamA$^S$. BamA$^S$ lacks POTRA domains 3, 4, and 5; however, the barrel is unaltered. This prevents BamA$^S$ from binding lipoproteins BamBCDE and forming an active folding complex upon completion of folding. The positions of the cysteine substitutions in each strand and loop used for BamA$^S$ disulfide crosslinking experiments are shown in yellow. Sequence identities of cysteine substitutions are listed in Table S4. (B) Ribbon model of the BamA β-barrel domain (PDB: 5D0O). Residues chosen for cysteine substitution for disulfide crosslinking experiments are shown as yellow sticks. Residues within β-strands that were chosen for cysteine substitution have outward-facing side chains and are positioned approximately in the middle of each β-strand. Residues within extracellular loops 1–5 and 7–8 that were chosen for cysteine substitution are positioned approximately in the middle of each loop. Residues within the large extracellular loop 6 that were chosen for cysteine substitution are distributed approximately evenly through the loop.

### Supplementary Tables

**Table S1.** *Escherichia coli* strains used in this study.

| Designation | Genotype or relevant description | Source or reference |
|---|---|---|
| NovaBlue | *endA1 hsdR17* ($r_{K12}^-$ $m_{K12}^+$) *supE44 thi-1 recA1 gyrA96 relA1 lac* F'[*proA*$^+$*B*$^+$*lacI*$^q$*Z*Δ*M15*::Tn*10*] | Novagen |
| DH5a *λpir* | F$^-$ Δ(*argF-lac*)*169* ϕ80d*lacZ58*(M15) Δ*phoA8 glnX44*(AS) *λ*$^-$ *deoR481 rfbC1 gyrA96*(NalR) *relA1 endA1 thiE1 hsdR17 λpir* | Laboratory collection (2) |
| MC4100 | F$^-$ [*araD139*]$_{B/r}$ Δ(*argF-lac*)*169 λ*$^-$ *e14*$^-$ *flhD5301* Δ(*fruK-yeiR*)*725*(*fruA25*) *relA1 rpsL150*(strR) *rbsR22* Δ(*fimB-fimE*)*632*(*::IS1*) *deoC1* | Coli Genetic Stock Center (CGSC: #6152) (3) |
| DEK1 | MC4100, *bamA-E470G* | (4) |
| DEK3 | MC4100, *bamA-E470K* | This study |
| DEK4 | MC4100, *lptD-ΔD330* | This study |
| DEK8 | MC4100, *lptD-ΔD330–D354* (*lptD4213*) | This study |
| DEK9 | MC4100, *lptD-N274I* | This study |
| DEK10 | MC4100, *lptD-ΔD330*, *N274I* | This study |
| DEK11 | MC4100, *lptD-ΔD330–D354*, *N274I* | This study |
| DEK31 | MC4100, *bamA-E470G*, *lptD-ΔD330* | This study |
| DEK32 | MC4100, *bamA-E470G*, *lptD-ΔD330–D354* | This study |
| DEK33 | MC4100, *bamA-E470K*, *lptD-ΔD330* | This study |
| BL21(DE3) | F$^-$ *ompT hsdS*$_B$ ($r_B^-$ $m_B^-$) *gal dcm* (DE3) | Novagen |
| NR1134 | MC4100, *ara*$^+$, Δ*lptD* [*lptD*-depletion strain], pACYC-*lptD* | Laboratory collection (5) |
| NR779 | MC4100, *degP::cam* | Laboratory collection (6) |

**Table S2.** Plasmids used in this study.

| Designation | Relevant description | Reference |
|---|---|---|
| pET23/42 | pET23a(+) containing part of the multiple cloning site of pET42a(+), T7 RNA polymerase-based expression system, pBR322 origin, $Crb^R$ | (7) |
| pET23/42-*lptD* | Encodes wild-type LptD | (8) |
| pET23/42-*lptD4213* | pET23/42-*lptD*-Δ*D330–D352* | (4) |
| pET23/42-*lptD*-Δ*D330–V334* | | This study |
| pET23/42-*lptD*-Δ*S335–Y339* | | This study |
| pET23/42-*lptD*-Δ*F343–Y347* | | This study |
| pET23/42-*lptD*-Δ*G348–D352* | | This study |
| pET23/42-*lptD-D330A–V334A* | | This study |
| pET23/42-*lptD-S335A–Y339A* | | This study |
| pET23/42-*lptD-F343A–Y347A* | | This study |
| pET23/42-*lptD-G348A–D352A* | | This study |
| pET23/42-*lptD*-Δ*D330* | | This study |
| pET23/42-*lptD*-Δ*Y331* | | This study |
| pET23/42-*lptD*-Δ*T332* | | This study |
| pET23/42-*lptD*-Δ*K333* | | This study |
| pET23/42-*lptD*-Δ*V334* | | This study |
| pET23/42-*lptD*-Δ*D330–Y331* | | This study |
| pET23/42-*lptD*-Δ*Y331–T332* | | This study |
| pET23/42-*lptD*-Δ*T332–K333* | | This study |
| pET23/42-*lptD*-Δ*K333–V334* | | This study |
| pET23/42-*lptD*-Δ*D330–T332* | | This study |
| pET23/42-*lptD*-Δ*Y331–K333* | | This study |
| pET23/42-*lptD*-Δ*T332–V334* | | This study |
| pET23/42-*lptD*-Δ*D330–K333* | | This study |
| pET23/42-*lptD*-Δ*Y331–V334* | | This study |
| pET23/42-*lptD*-Δ*D330*/Δ*K333–D352* | | This study |
| pET23/42-*lptD*-Δ*K333–D352* | | This study |
| pET23/42-*lptD*-Δ*K375–V379* | | This study |
| pSup-BpaRS-6TRN | Encodes an archaeal aminoacyl tRNA synthetase and an amber suppressor tRNA that facilitates incorporation of the photoreactive, unnatural amino acid *p*- | (9) |

| | | |
|---|---|---|
| | benzoyl-L-phenylalanine (*p*Bpa) into a protein of interest, $Cam^R$ | |
| pZS21-*lptD*-8×His | Encodes a variant of LptD containing an octahistidine tag at the C-terminal region, tetracycline-inducible expression system, pSC101 origin, $Kan^R$ | (4) |
| pZS21-*lptD*-Δ*D330, E435X*-8×His | | This study |
| pZS21-*lptD*-Δ*D330, S439X*-8×His | | This study |
| pZS21-*lptD*-Δ*D330, Q441X*-8×His | | This study |
| pZS21-*lptD*-Δ*D330, Y468X*-8×His | | This study |
| pZS21-*lptD*-Δ*D330, E470X*-8×His | | This study |
| pZS21-*lptD*-Δ*D330, L471X*-8×His | | This study |
| pZS21-*lptD*-Δ*D330, R488X*-8×His | | This study |
| pZS21-*lptD*-Δ*D330, F490X*-8×His | | This study |
| pZS21-*lptD*-Δ*D330, N492X*-8×His | | This study |
| pZS21-*lptD*-Δ*D330, F494X*-8×His | | This study |
| pZS21-*lptD*-Δ*D330, L501X*-8×His | | This study |
| pZS21-*lptD*-Δ*D330, T511X*-8×His | | This study |
| pZS21-*lptD*-Δ*D330, G528X*-8×His | | This study |
| pZS21-*lptD*-Δ*D330, G530X*-8×His | | This study |
| pZS21-*lptD*-Δ*D330, D569X*-8×His | | This study |
| pZS21-*lptD*-Δ*D330, K610*X-8×His | | This study |
| pZS21-*lptD*-Δ*D330, D614X*-8×His | | This study |
| pZS21-*lptD*-Δ*D330, T615X*-8×His | | This study |
| pZS21-6×His-*bamA* | Encodes a variant of BamA containing an hexahistidine tag at the N-terminal region, tetracycline-inducible expression system, pSC101 origin, $Kan^R$ | (10) |
| pZS21-6×His-*bamA-Q466X* | | This study |
| pZS21-6×His-*bamA-Y468X* | | This study |
| pZS21-6×His-*bamA-S472X* | | This study |
| pZS21-6×His-*bamA-T474X* | | This study |
| pZS21-6×His-*bamA-Y468C* | | This study |
| pZS21-6×His-*bamA-D512C* | | This study |
| pZS21-6×His-*bamA-E470G, D512C* | | This study |
| pTrc99A-*lptD*-3×FLAG | Encodes a variant of LptD containing a 3×FLAG tag at the C-terminal region, IPTG-inducible expression system, pBR322 origin, $Crb^R$ | (4) |

| | |
|---|---|
| pTrc99A-*lptD-Y235C*-3×FLAG | This study |
| pTrc99A-*lptD-F243C*-3×FLAG | This study |
| pTrc99A-*lptD-I259C*-3×FLAG | This study |
| pTrc99A-*lptD-W272C*-3×FLAG | This study |
| pTrc99A-*lptD-M287C*-3×FLAG | This study |
| pTrc99A-*lptD-W315C*-3×FLAG | This study |
| pTrc99A-*lptD-F327C*-3×FLAG | This study |
| pTrc99A-*lptD-F359C*-3×FLAG | This study |
| pTrc99A-*lptD-V372C*-3×FLAG | This study |
| pTrc99A-*lptD-Y398C*-3×FLAG | This study |
| pTrc99A-*lptD-I410C*-3×FLAG | This study |
| pTrc99A-*lptD-L432C*-3×FLAG | This study |
| pTrc99A-*lptD-L453C*-3×FLAG | This study |
| pTrc99A-*lptD-V487C*-3×FLAG | This study |
| pTrc99A-*lptD-P510C*-3×FLAG | This study |
| pTrc99A-*lptD-V562C*-3×FLAG | This study |
| pTrc99A-*lptD-V579C*-3×FLAG | This study |
| pTrc99A-*lptD-W608C*-3×FLAG | This study |
| pTrc99A-*lptD-I626C*-3×FLAG | This study |
| pTrc99A-*lptD-S640C*-3×FLAG | This study |
| pTrc99A-*lptD-Y656C*-3×FLAG | This study |
| pTrc99A-*lptD-A687C*-3×FLAG | This study |
| pTrc99A-*lptD-Y703C*-3×FLAG | This study |
| pTrc99A-*lptD-S715C*-3×FLAG | This study |
| pTrc99A-*lptD-R734C*-3×FLAG | This study |
| pTrc99A-*lptD-N750C*-3×FLAG | This study |
| pTrc99A-*lptD-V635C*-3×FLAG | This study |
| pTrc99A-*lptD-Y663C*-3×FLAG | This study |
| pTrc99A-*lptD-T238C*-3×FLAG | This study |
| pTrc99A-*lptD-R267C*-3×FLAG | This study |
| pTrc99A-*lptD-N304C*-3×FLAG | This study |
| pTrc99A-*lptD-D344C*-3×FLAG | This study |
| pTrc99A-*lptD-Q383C*-3×FLAG | This study |
| pTrc99A-*lptD-D422C*-3×FLAG | This study |
| pTrc99A-*lptD-T472C*-3×FLAG | This study |
| pTrc99A-*lptD-D536C*-3×FLAG | This study |
| pTrc99A-*lptD-N594C*-3×FLAG | This study |

| | |
|---|---|
| pTrc99A-*lptD-R631C*-3×FLAG | This study |
| pTrc99A-*lptD-K670C*-3×FLAG | This study |
| pTrc99A-*lptD-N708C*-3×FLAG | This study |
| pTrc99A-*lptD-D742C*-3×FLAG | This study |
| pTrc99A-*lptD4213-F243C*-3×FLAG | This study |
| pTrc99A-*lptD4213-I259C*-3×FLAG | This study |
| pTrc99A-*lptD4213-W272C*-3×FLAG | This study |
| pTrc99A-*lptD4213-M287C*-3×FLAG | This study |
| pTrc99A-*lptD4213-W315C*-3×FLAG | This study |
| pTrc99A-*lptD4213-F327C*-3×FLAG | This study |
| pTrc99A-*lptD4213-F359C*-3×FLAG | This study |
| pTrc99A-*lptD4213-L453C*-3×FLAG | This study |
| pTrc99A-*lptD4213-V487C*-3×FLAG | This study |
| pTrc99A-*lptD4213-P510C*-3×FLAG | This study |
| pTrc99A-*lptD4213-V562C*-3×FLAG | This study |
| pTrc99A-*lptD4213-V579C*-3×FLAG | This study |
| pTrc99A-*lptD4213-W608C*-3×FLAG | This study |
| pTrc99A-*lptD4213-I626C*-3×FLAG | This study |
| pTrc99A-*lptD4213-S640C*-3×FLAG | This study |
| pTrc99A-*lptD4213-Y656C*-3×FLAG | This study |
| pTrc99A-*lptD4213-A687C*-3×FLAG | This study |
| pTrc99A-*lptD4213-Y703C*-3×FLAG | This study |
| pTrc99A-*lptD4213-S715C*-3×FLAG | This study |
| pTrc99A-*lptD4213-R734C*-3×FLAG | This study |
| pTrc99A-*lptD4213-N750C*-3×FLAG | This study |
| pTrc99A-*lptD-ΔD330, F243C*-3×FLAG | This study |
| pTrc99A-*lptD-ΔD330, I259C*-3×FLAG | This study |
| pTrc99A-*lptD-ΔD330, W272C*-3×FLAG | This study |
| pTrc99A-*lptD-ΔD330, M287C*-3×FLAG | This study |
| pTrc99A-*lptD-ΔD330, W315C*-3×FLAG | This study |

| | |
|---|---|
| pTrc99A-*lptD-ΔD330, F327C*-3×FLAG | This study |
| pTrc99A-*lptD-ΔD330, F359C*-3×FLAG | This study |
| pTrc99A-*lptD-ΔD330, L453C*-3×FLAG | This study |
| pTrc99A-*lptD-ΔD330, V487C*-3×FLAG | This study |
| pTrc99A-*lptD-ΔD330, P510C*-3×FLAG | This study |
| pTrc99A-*lptD-ΔD330, V562C*-3×FLAG | This study |
| pTrc99A-*lptD-ΔD330, V579C*-3×FLAG | This study |
| pTrc99A-*lptD-ΔD330, W608C*-3×FLAG | This study |
| pTrc99A-*lptD-ΔD330, I626C*-3×FLAG | This study |
| pTrc99A-*lptD-ΔD330, S640C*-3×FLAG | This study |
| pTrc99A-*lptD-ΔD330, Y656C*-3×FLAG | This study |
| pTrc99A-*lptD-ΔD330, A687C*-3×FLAG | This study |
| pTrc99A-*lptD-ΔD330, Y703C*-3×FLAG | This study |
| pTrc99A-*lptD-ΔD330, S715C*-3×FLAG | This study |
| pTrc99A-*lptD-ΔD330, R734C*-3×FLAG | This study |
| pTrc99A-*lptD-ΔD330, N750C*-3×FLAG | This study |
| pTrc99A-*lptD-ΔD330, T238C*-3×FLAG | This study |
| pTrc99A-*lptD-ΔD330, R267C*-3×FLAG | This study |
| pTrc99A-*lptD-ΔD330, N304C*-3×FLAG | This study |
| pTrc99A-*lptD-ΔD330, D344C*-3×FLAG | This study |
| pTrc99A-*lptD-ΔD330, Q383C*-3×FLAG | This study |

| | | |
|---|---|---|
| pTrc99A-*lptD-ΔD330, D422C*-3×FLAG | | This study |
| pTrc99A-*lptD-ΔD330, T472C*-3×FLAG | | This study |
| pTrc99A-*lptD-ΔD330, D536C*-3×FLAG | | This study |
| pTrc99A-*lptD-ΔD330, N594C*-3×FLAG | | This study |
| pTrc99A-*lptD-ΔD330, R631C*-3×FLAG | | This study |
| pTrc99A-*lptD-ΔD330, K670C*-3×FLAG | | This study |
| pTrc99A-*lptD-ΔD330, N708C*-3×FLAG | | This study |
| pTrc99A-*lptD-ΔD330, D742C*-3×FLAG | | This study |
| pTrc99A-3×FLAG-*bamA-ΔG172–R421* | Encodes a variant of BamA containing a 3×FLAG tag at the N-terminal region and lacking the periplasmic polypeptide transport-associated (POTRA) domains 3–5 (amino acids G172–R421), IPTG-inducible expression system, pBR322 origin, $Crb^R$ | (11) |
| pTrc99A-3×FLAG-*bamA-ΔG172–R421, F428C* | | This study |
| pTrc99A-3×FLAG-*bamA-ΔG172–R421, A442C* | | This study |
| pTrc99A-3×FLAG-*bamA-ΔG172–R421, I458C* | | This study |
| pTrc99A-3×FLAG-*bamA-ΔG172–R421, L471C* | | This study |
| pTrc99A-3×FLAG-*bamA-ΔG172–R421, L489C* | | This study |
| pTrc99A-3×FLAG-*bamA-ΔG172–R421, T511C* | | This study |
| pTrc99A-3×FLAG-*bamA-ΔG172–R421, L529C* | | This study |
| pTrc99A-3×FLAG-*bamA-ΔG172–R421, F572C* | | This study |
| pTrc99A-3×FLAG-*bamA-ΔG172–R421, L595C* | | This study |
| pTrc99A-3×FLAG-*bamA-ΔG172–R421, L613C* | | This study |
| pTrc99A-3×FLAG-*bamA-ΔG172–R421, W635C* | | This study |

| | | |
|---|---|---|
| pTrc99A-3×FLAG-*bamA-ΔG172–R421, A714C* | | This study |
| pTrc99A-3×FLAG-*bamA-ΔG172–R421, W739C* | | This study |
| pTrc99A-3×FLAG-*bamA-ΔG172–R421, I772C* | | This study |
| pTrc99A-3×FLAG-*bamA-ΔG172–R421, F785C* | | This study |
| pTrc99A-3×FLAG-*bamA-ΔG172–R421, F804C* | | This study |
| pTrc99A-3×FLAG-*bamA-ΔG172–R421, E435C* | | This study |
| pTrc99A-3×FLAG-*bamA-ΔG172–R421, D464C* | | This study |
| pTrc99A-3×FLAG-*bamA-ΔG172–R421, D500C* | | This study |
| pTrc99A-3×FLAG-*bamA-ΔG172–R421, R547C* | | This study |
| pTrc99A-3×FLAG-*bamA-ΔG172–R421, S604C* | | This study |
| pTrc99A-3×FLAG-*bamA-ΔG172–R421, E650C* | | This study |
| pTrc99A-3×FLAG-*bamA-ΔG172–R421, Q664C* | | This study |
| pTrc99A-3×FLAG-*bamA-ΔG172–R421, Y674C* | | This study |
| pTrc99A-3×FLAG-*bamA-ΔG172–R421, D685C* | | This study |
| pTrc99A-3×FLAG-*bamA-ΔG172–R421, D694C* | | This study |
| pTrc99A-3×FLAG-*bamA-ΔG172–R421, D704C* | | This study |
| pTrc99A-3×FLAG-*bamA-ΔG172–R421, S755C* | | This study |
| pTrc99A-3×FLAG-*bamA-ΔG172–R421, D795C* | | This study |
| pTrc99A-*lptD-Y721D*-3×FLAG | | (5) |
| pTrc99A-8×His-*bamA-Y468Bpa-bamBCDE* | Encodes a variant of BamA containing a octahistidine tag at the N-terminal region and a replacement of residue Y468 for an amber codon (TAG), the four lipoproteins (BamBCDE), IPTG-inducible expression system, pBR322 origin, $Crb^R$ | Modified from (12) |

| | | |
|---|---|---|
| pCDFDuet1-*lptE*-2×Strep-tag-*lptD∆D330* | | This study |
| pDS132 | Suicide vector for allelic exchange, R6K origin, $Chl^R$ | (13) |
| pDS132-*bamA* | Encodes a wild type BamA | (4) |
| pDS132-*bamA-E470G* | | (4) |
| pDS132-*bamA-E470K* | | This study |
| pDS132-*lptD* | Encodes a wild type LptD | This study |
| pDS132-*lptD4213* | | This study |
| pDS132-*lptD-∆D330* | | This study |
| pDS132-*lptD-N274I* | | This study |
| pDS132-*lptD4213-N274I* | | This study |
| pDS132-*lptD-∆D330, N274I* | | This study |

**Table S3.** Oligonucleotide primers used in this study.

| **Primer**[a] | **Sequence (5'–3')** |
|---|---|
| *Sequencing primers*[b] | |
| *lptD554*-F | gcgaaattattcatgaccgc |
| *lptD1157*-F | gtagctactcggcagagcc |
| *lptD1751*-F | atttcacggagtctcgcact |
| *lptD868*-R | ccagttccatcaagccagc |
| *lptE142*-F | agccgtgcggtgcgtaaccag |
| *lptE441*-R | catctcttttacgatcatgtc |
| *lptD330*-upstream-F | ctaacactacgccccaaaag |
| *lptD334*-downstream-R | ctcatcggagattttcactc |
| *bamA474*-F | gccgcgcaaccgtgttgacc |
| *bamA1179*-F | cttctttgaaactgtcgatacc |
| *bamA717*-R | gttgaaacgggcataacc |
| *bamA1875*-F | caaatgggttgttctggg |
| *bamB484*-F | gtgttaatccacaccagt |
| *bamB688*-R | gctgccaaatcatctggc |
| *bamC417*-F | gacactgaccaccgattg |
| *bamC617*-R | gaaataacgttcatcatc |
| *bamD269*-F | ctgccatcgatcgtttta |
| *bamD470*-R | ctgttcggatagccgcgc |
| *bamE67*-F | ctggagcgagtggtttaccg |
| *bamE270*-R | cgtttgctgagttacaccttc |
| *Mutagenesis primer*[c] | |
| *lptD*-wt-F | atggatcaggtgtggcgtttcaacgtcgactacaccaaggtcagcgatcctagctac |
| *lptD*-wt-R | gtagctaggatcgctgaccttggtgtagtcgacgttgaaacgccacacctgatccat |
| *lptD*-ΔD330-D352-F | gtcatggatcaggtgtggcgtttcaacgtcggctacgcaacgcaaaaattcagcgttggc |
| *lptD*-ΔD330-D352-R | gccaacgctgaatttttgcgttgcgtagccgacgttgaaacgccacacctgatccatgac |
| *lptD*-ΔD330-V334-F | ggggtcatggatcaggtgtggcgtttcaacgtcagcgatcctagctacttcaatgatttcg |
| *lptD*-ΔD330-V334-R | cgaaatcattgaagtagctaggatcgctgacgttgaaacgccacacctgatccatgacccc |
| *lptD*-ΔS335-Y339-F | gtgtggcgtttcaacgtcgactacaccaaggtcttcaatgatttcgataacaagtacggttcc |
| *lptD*-ΔS335-Y339-R | ggaaccgtacttgttatcgaaatcattgaagaccttggtgtagtcgacgttgaaacgccacac |
| *lptD*-ΔF343-Y347-F | ccaaggtcagcgatcctagctacttcaatgatggttccagtactgacggctacgcaacgc |
| *lptD*-ΔF343-Y347-R | gcgttgcgtagccgtcagtactggaaccatcattgaagtagctaggatcgctgaccttgg |
| *lptD*-ΔG348-D352-F | ctagctacttcaatgatttcgataacaagtacggctacgcaacgcaaaaattcagcgttggc |
| *lptD*-ΔG348-D352-R | gccaacgctgaatttttgcgttgcgtagccgtacttgttatcgaaatcattgaagtagctag |
| *lptD*-D330A-V334A-F | gatcaggtgtggcgtttcaacgtc**gccgcggcagctgcc**agcgatcctagctacttcaatg |
| *lptD*-D330A-V334A-R | cattgaagtagctaggatcgct**ggcagctgccgcggc**gacgttgaaacgccacacctgatc |
| *lptD*-S335A-Y339A-F | caacgtcgactacaccaaggtc**gccgcggcagctgcc**ttcaatgatttcgataacaagtac |
| *lptD*-S335A-Y339A-R | gtacttgttatcgaaatcattgaa**ggcagctgccgcggc**gaccttggtgtagtcgacgttg |
| *lptD*-F343A-Y347A-F | gcgatcctagctacttcaatgat**gccgcggcagctgcc**ggttccagtactgacggctacg |
| *lptD*-F343A-Y347A-R | cgtagccgtcagtactggaacc**ggcagctgccgcggc**atcattgaagtagctaggatcgc |
| *lptD*-G348A-D352A-F | caatgatttcgataacaagtac**gccgcggcagctgcc**ggctacgcaacgcaaaaattcagcg |
| *lptD*-G348A-D352A-R | cgctgaatttttgcgttgcgtagcc**ggcagctgccgcggc**gtacttgttatcgaaatcattg |
| *lptD*-ΔD330-F | gtcatggatcaggtgtggcgtttcaacgtctacaccaaggtcagcgatcctagctacttc |
| *lptD*-ΔD330-R | gaagtagctaggatcgctgaccttggtgtagacgttgaaacgccacacctgatccatgac |
| *lptD*-ΔY331-F | ggatcaggtgtggcgtttcaacgtcgacaccaaggtcagcgatcctagctacttcaatg |
| *lptD*-ΔY331-R | cattgaagtagctaggatcgctgaccttggtgtcgacgttgaaacgccacacctgatcc |
| *lptD*-ΔT332-F | gatcaggtgtggcgtttcaacgtcgactacaaggtcagcgatcctagctacttcaatg |
| *lptD*-ΔT332-R | cattgaagtagctaggatcgctgaccttgtagtcgacgttgaaacgccacacctgatc |
| *lptD*-ΔK333-F | caggtgtggcgtttcaacgtcgactacaccgtcagcgatcctagctacttcaatgatttc |
| *lptD*-ΔK333-R | gaaatcattgaagtagctaggatcgctgacggtgtagtcgacgttgaaacgccacacctg |
| *lptD*-ΔV334-F | gtgtggcgtttcaacgtcgactacaccaagagcgatcctagctacttcaatgatttcg |

| | |
|---|---|
| *lptD*-ΔV334-R | cgaaatcattgaagtagctaggatcgctcttggtgtagtcgacgttgaaacgccacac |
| *lptD*-ΔD330-Y331-F | catggatcaggtgtggcgtttcaacgtcaccaaggtcagcgatcctagctacttcaatg |
| *lptD*-ΔD330-Y331-R | cattgaagtagctaggatcgctgaccttggtgacgttgaaacgccacacctgatccatg |
| *lptD*-ΔY331-T332-F | catggatcaggtgtggcgtttcaacgtcgacaaggtcagcgatcctagctacttcaatgatttc |
| *lptD*-ΔY331-T332-R | gaaatcattgaagtagctaggatcgctgaccttgtcgacgttgaaacgccacacctgatccatg |
| *lptD*-ΔT332-K333-F | gatcaggtgtggcgtttcaacgtcgactacgtcagcgatcctagctacttcaatgatttc |
| *lptD*-ΔT332-K333-R | gaaatcattgaagtagctaggatcgctgacgtagtcgacgttgaaacgccacacctgatc |
| *lptD*-ΔK333-V334-F | caggtgtggcgtttcaacgtcgactacaccagcgatcctagctacttcaatgatttcg |
| *lptD*-ΔK333-V334-R | cgaaatcattgaagtagctaggatcgctggtgtagtcgacgttgaaacgccacacctg |
| *lptD*-ΔD330-T332-F | catggatcaggtgtggcgtttcaacgtcaaggtcagcgatcctagctacttcaatgatttc |
| *lptD*-ΔD330-T332-R | gaaatcattgaagtagctaggatcgctgaccttgacgttgaaacgccacacctgatccatg |
| *lptD*-ΔY331-K333-F | gtcatggatcaggtgtggcgtttcaacgtcgacgtcagcgatcctagctacttcaatgatttc |
| *lptD*-ΔY331-K333-R | gaaatcattgaagtagctaggatcgctgacgtcgacgttgaaacgccacacctgatccatgac |
| *lptD*-ΔT332-V334-F | gatcaggtgtggcgtttcaacgtcgactacagcgatcctagctacttcaatgatttcg |
| *lptD*-ΔT332-V334-R | cgaaatcattgaagtagctaggatcgctgtagtcgacgttgaaacgccacacctgatc |
| *lptD*-ΔD330-K333-F | gtcatggatcaggtgtggcgtttcaacgtcgtcagcgatcctagctacttcaatgatttc |
| *lptD*-ΔD330-K333-R | gaaatcattgaagtagctaggatcgctgacgacgttgaaacgccacacctgatccatgac |
| *lptD*-ΔY331-V334-F | ggatcaggtgtggcgtttcaacgtcgacagcgatcctagctacttcaatgatttcg |
| *lptD*-ΔY331-V334-R | cgaaatcattgaagtagctaggatcgctgtcgacgttgaaacgccacacctgatcc |
| *lptD*-ΔD330/ΔK333-D352-F | gtcatggatcaggtgtggcgtttcaacgtctacaccggctacgcaacgcaaaaattcagcgttggc |
| *lptD*-ΔD330/ΔK333-D352-R | gccaacgctgaatttttgcgttgcgtagccggtgtagacgttgaaacgccacacctgatccatgac |
| *lptD*-ΔK333-D352-F | caggtgtggcgtttcaacgtcgactacaccggctacgcaacgcaaaaattcagcgttggc |
| *lptD*-ΔK333-D352-R | gccaacgctgaatttttgcgttgcgtagccggtgtagtcgacgttgaaacgccacacctg |
| *lptD*-ΔK375-V379-F | gtgcaaaacttcaatgccaccgtttcaaccttcagcgaacagaacaccagtagctactcg |
| *lptD*-ΔK375-V379-R | cgagtagctactggtgttctgttcgctgaaggttgaaacggtggcattgaagttttgcac |
| *lptD*-Y235C-F | ctctggtttcttgatcccgaacgccaag**tgc**accaccaccaactactttgagttctac |
| *lptD*-Y235C-R | gtagaactcaaagtagttggtggtggt**gca**cttggcgttcgggatcaagaaaccagag |
| *lptD*-F243C-F | gtacaccaccaccaactactttgag**tgc**tacctgccatattactggaacatcgcgcc |
| *lptD*-F243C-R | ggcgcgatgttccagtaatatggcaggta**gca**ctcaaagtagttggtggtggtgtac |
| *lptD*-I259C-F | gaacatcgcgccaaatatggatgccacc**tgc**acgccgcattatatgcatcgtcgtggc |
| *lptD*-I259C-R | gccacgacgatgcatataatgcggcgt**gca**ggtggcatccatatttggcgcgatgttc |
| *lptD*-W272C-F | cattatatgcatcgtcgtggcaacatcatg**tgt**gagaacgaattccgctacctctcccag |
| *lptD*-W272C-R | ctgggagaggtagcggaattcgttctc**aca**catgatgttgccacgacgatgcatataatg |
| *lptD*-M287C-F | ctacctctcccaggcgggcgctggcttg**tgc**gaactggactatctgccttcagataaag |
| *lptD*-M287C-R | ctttatctgaaggcagatagtccagttc**gca**caagccagcgcccgcctgggagaggtag |
| *lptD*-W315C-F | gacagttcacgtcgttggttattctac**tgt**aaccactccggggtcatggatcaggtg |
| *lptD*-W315C-R | cacctgatccatgaccccggagtggtt**aca**gtagaataaccaacgacgtgaactgtc |
| *lptD*-F327C-F | ggtgtggcgt**tgc**aacgtcgact |
| *lptD*-F327C-R | agtcgacgtt**gca**acgccacacc |
| *lptD*-F327C-DD330-F | ggtgtggcgt**tgc**aacgtctaca |
| *lptD*-F327C-DD330-R | tgtagacgtt**gca**acgccacacc |
| *lptD*-F327C-DD330-D352-F | ggtgtggcgt**tgc**aacgtcggct |
| *lptD*-F327C-DD330-D352-R | agccgacgtt**gca**acgccacacc |
| *lptD*-Y331C-F | gatcaggtgtggcgtttcaacgtcgac**tgc**accaaggtcagcgatcctagctacttc |
| *lptD*-Y331C-R | gaagtagctaggatcgctgaccttggt**gca**gtcgacgttgaaacgccacacctgatc |
| *lptD*-DD330-Y331C-F | atggatcaggtgtggcgtttcaacgtc**tgc**accaaggtcagcgatcctagctacttc |
| *lptD*-DD330-Y331C-R | gaagtagctaggatcgctgaccttggt**gca**gacgttgaaacgccacacctgatccat |
| *lptD*-F359C-F | cagtactgacggctacgcaacgcaaaaa**tgc**agcgttggctatgcggtgcaaaacttc |
| *lptD*-F359C-R | gaagttttgcaccgcatagccaacgct**gca**tttttgcgttgcgtagccgtcagtactg |
| *lptD*-V372C-F | ctatgcggtgcaaaacttcaatgccacc**tgt**tcaaccaagcagttccaggttttcagc |
| *lptD*-V372C-R | gctgaaaacctggaactgcttggttga**aca**ggtggcattgaagttttgcaccgcatag |
| *lptD*-Y398C-F | ctcggcagagccgcagttagacgttaat**tgc**taccagaatgatgttggtccgtttgatac |
| *lptD*-Y398C-R | gtatcaaacggaccaacatcattctggta**gca**attaacgtctaactgcggctctgccgag |
| *lptD*-I410C-F | tgatacgcgt**tgc**tacggccagg |

| | |
|---|---|
| *lptD*-I410C-R | cctggccgta**gca**acgcgtatca |
| *lptD*-L432C-F | ccgtgttcac**tgt**gaaccgacca |
| *lptD*-L432C-R | tggtcggttc**aca**gtgaacacgg |
| *lptD*-L453C-F | cgaagcgaag**tgc**ctggcaaccc |
| *lptD*-L453C-R | gggttgccag**gca**cttcgcttcg |
| *lptD*-V487C-F | gcaattcaaa**tgc**gacggcaaaa |
| *lptD*-V487C-R | ttttgccgtc**gca**tttgaattgc |
| *lptD*-P510C-F | aacgctggaa**tgc**cgcgcgcagt |
| *lptD*-P510C-R | actgcgcgcg**gca**ttccagcgtt |
| *lptD*-V562C-F | gacgaccggt**tgc**acatctcgca |
| *lptD*-V562C-R | tgcgagatgt**gca**accggtcgtc |
| *lptD*-V579C-F | taatatttcc**tgc**ggtcaaatct |
| *lptD*-V579C-R | agatttgacc**gca**ggaaatatta |
| *lptD*-W608C-F | aatgacgacaaaacgggttcactggtg**tgc**gcaggcgatacttactggcgtatctcc |
| *lptD*-W608C-R | ggagatacgccagtaagtatcgcctgc**gca**caccagtgaacccgttttgtcgtcatt |
| *lptD*-I626C-F | gcgtggcggg**tgc**cagtacgata |
| *lptD*-I626C-R | tatcgtactg**gca**cccgccacgc |
| *lptD*-S640C-F | cgtctggataacgtagcgaccagtaac**tgc**agcattgaataccgtcgggatgaagac |
| *lptD*-S640C-R | gtcttcatcccgacggtattcaatgct**gca**gttactggtcgctacgttatccagacg |
| *lptD*-Y656C-F | acagctgaat**tgc**cgttacgcca |
| *lptD*-Y656C-R | tggcgtaacg**gca**attcagctgt |
| *lptD*-A687C-F | tataagaatggtatttcgcaggtaggt**tgc**gtcgccagctggccaattgccgatcgt |
| *lptD*-A687C-R | acgatcggcaattggccagctggcgac**gca**acctacctgcgaaataccattcttata |
| *lptD*-Y703C-F | tgttggggcc**tgc**tactacgaca |
| *lptD*-Y703C-R | tgtcgtagta**gca**ggccccaaca |
| *lptD*-S715C-F | gacaccaatgctaacaagcaagccgac**tgc**atgttaggtgtgcaatacagctcctgc |
| *lptD*-S715C-R | gcaggagctgtattgcacacctaacat**gca**gtcggcttgcttgttagcattggtgtc |
| *lptD*-R734C-F | cggttacgag**tgc**aagctgaacg |
| *lptD*-R734C-R | cgttcagctt**gca**ctcgtaaccg |
| *lptD*-N750C-F | gtatatgac**tgc**gcaatcggc |
| *lptD*-N750C-R | gccgattgc**gca**gtcatatac |
| *lptD*-T238C-F | gtacaccacc**tgc**aactactttg |
| *lptD*-T238C-R | caaagtagtt**gca**ggtggtgtac |
| *lptD*-R267C-F | tatgcatcgt**tgc**ggcaacatca |
| *lptD*-R267C-R | tgatgttgcc**gca**acgatgcata |
| *lptD*-N304C-F | tgaacacccg**tgc**gatgacagtt |
| *lptD*-N304C-R | aactgtcatc**gca**cgggtgttca |
| *lptD*-D344C-F | caatgatttc**tgc**aacaagtacg |
| *lptD*-D344C-R | cgtacttgtt**gca**gaaatcattg |
| *lptD*-Q383C-F | tttcagcgaa**tgc**aacaccagta |
| *lptD*-Q383C-R | tactggtgtt**gca**ttcgctgaaa |
| *lptD*-D422C-F | taacaccaga**tgc**gacatgcctg |
| *lptD*-D422C-R | caggcatgtc**gca**tctggtgtta |
| *lptD*-T472C-F | cagaaacacg**tgc**aagctggacg |
| *lptD*-T472C-R | cgtccagctt**gca**cgtgtttctg |
| *lptD*-D536C-F | gctgcaatct**tgc**tactctggcc |
| *lptD*-D536C-R | ggccagagta**gca**agattgcagc |
| *lptD*-N594C-F | tggcgatgac**tgc**ataacatggg |
| *lptD*-N594C-R | cccatgttat**gca**gtcatcgcca |
| *lptD*-R631C-F | gtacgataca**tgc**ctggataacg |
| *lptD*-R631C-R | cgttatccag**gca**tgtatcgtac |
| *lptD*-K670C-F | tacgctgcct**tgc**tactattcca |
| *lptD*-K670C-R | tggaatagta**gca**aggcagcgta |
| *lptD*-N708C-F | ctacgacacc**tgc**gctaacaagc |
| *lptD*-N708C-R | gcttgttagc**gca**ggtgtcgtag |
| *lptD*-D742C-F | ttgggataac**tgc**aaacaacatg |

| | |
|---|---|
| *lptD*-D742C-R | catgttgttt**gca**gttatcccaa |
| *bamA*-Q466*p*BPA-F | ggtatcaacgggaccaaaaacgattac**tag**acctatgctgaactgtcggtaaccaac |
| *bamA*-Q466*p*BPA-R | gttggttaccgacagttcagcataggt**cta**gtaatcgtttttggtcccgttgatacc |
| *bamA*-Y468*p*BPA-F | caacgggaccaaaaacgattaccagacc**tag**gctgaactgtcggtaaccaacccgtac |
| *bamA*-Y468*p*BPA-R | gtacgggttggttaccgacagttcagc**cta**ggtctggtaatcgtttttggtcccgttg |
| *bamA*-S472*p*BPA-F | cgattaccagacctatgctgaactg**tag**gtaaccaacccgtacttcaccgtagatgg |
| *bamA*-S472*p*BPA-R | ccatctacggtgaagtacgggttggttac**cta**cagttcagcataggtctggtaatcg |
| *bamA*-T474*p*BPA-F | ccagacctatgctgaactgtcggta**tag**aacccgtacttcaccgtagatggcgtaag |
| *bamA*-T474*p*BPA-R | cttacgccatctacggtgaagtacgggtt**cta**taccgacagttcagcataggtctgg |
| *bamA*-Q441C-F | cgtgagcttc**tgc**gctggtgtgc |
| *bamA*-Q441C-R | gcacaccagc**gca**gaagctcacg |
| *bamA*-Y468C-F | caacgggaccaaaaacgattaccagacc**tgc**gctgaactgtcggtaaccaacccgtac |
| *bamA*-Y468C-R | gtacgggttggttaccgacagttcagc**gca**ggtctggtaatcgtttttggtcccgttg |
| *bamA*-D512C-F | ttatggtaca**tgc**gtgacgttgg |
| *bamA*-D512C-R | ccaacgtcac**gca**tgtaccataa |
| *bamA*-E470G-F | accaaaaacgattaccagacctatgct**gga**ctgtcggtaaccaacccgtacttcacc |
| *bamA*-E470G-R | ggtgaagtacgggttggttaccgacag**tcc**agcataggtctggtaatcgtttttggt |
| *bamA*-Y468C-E470G-F | gggaccaaaaacgattaccagacc**tgc**gct**ggc**ctgtcggtaaccaacccgtacttc |
| *bamA*-Y468C-E470G-R | gaagtacgggttggttaccgacag**gcc**agc**gca**ggtctggtaatcgtttttggtccc |
| *Cloning primers*[d] | |
| pDS132-linear-F | ggatccgtcgacctgcagatccgtcgacct |
| pDS132-linear-R | ggatcccaagcttcttctagaggtaccgca |
| *lptD*-pDS132-F | **aggtcgacggatctgcaggtcgacggatcc**atgaaaaaacgtatccccactctc |
| *lptD*-pDS132-R | **tgcggtacctctagaagaagcttgggatcc**tcacaaagtgttttgatacggcaga |

[a] Oligonucleotide primers were from Eton Bioscience or Genewiz.

[b] Sequencing primers names correspond to the position and gene they target.

[c] Mutagenesis primers names correspond to the gene and codon where the mutation (substitutions or deletion) is located. For codon substitutions (highlighted in bold), we replaced the original codon (or a segment of codons), encoding the amino acid indicated by the one-letter code that precedes the codon number, with a codon (or a segment of codons) that encodes the amino acid indicated by the one-letter code that follows the codon number. To introduce the unnatural amino acid *p*-benzoyl-L-phenylalanine (*p*BPA), we replaced the original codon with an amber stop codon (highlighted in red)

[d] Cloning primers names correspond to the gene and plasmid vector used in the assembly reaction. Complementary overlapping region is in bold and underlined.

**Table S4.** Sequence identity of substrates used in this study.

| Substrate | Mutation position | Region of the substrate |
|---|---|---|
| LptD | Y235C | β-strand 1 |
| | F243C | β-strand 2 |
| | I259C | β-strand 3 |
| | W272C | β-strand 4 |
| | M287C | β-strand 5 |
| | W315C | β-strand 6 |
| | F327C | β-strand 7 |
| | F359C | β-strand 8 |
| | V372C | β-strand 9 |
| | Y398C | β-strand 10 |
| | I410C | β-strand 11 |
| | L432C | β-strand 12 |
| | L453C | β-strand 13 |
| | V487C | β-strand 14 |
| | P510C | β-strand 15 |
| | V562C | β-strand 16 |
| | V579C | β-strand 17 |
| | W608C | β-strand 18 |
| | I626C | β-strand 19 |
| | S640C | β-strand 20 |
| | Y656C | β-strand 21 |
| | A687C | β-strand 22 |
| | Y703C | β-strand 23 |
| | S715C | β-strand 24 |
| | R734C | β-strand 25 |
| | N759C | β-strand 26 |
| | T238C | extracellular loop 1 |
| | R267C | extracellular loop 2 |
| | N304C | extracellular loop 3 |
| | D344C | extracellular loop 4 |
| | Q383C | extracellular loop 5 |
| | D422C | extracellular loop 6 |
| | T472C | extracellular loop 7 |
| | D536C | extracellular loop 8 |
| | N594C | extracellular loop 9 |
| | R631C | extracellular loop 10 |
| | K670C | extracellular loop 11 |
| | N708C | extracellular loop 12 |
| | D742C | extracellular loop 13 |
| $BamA^S$ | F428C | β-strand 1 |
| | A442C | β-strand 2 |
| | I458C | β-strand 3 |
| | L471C | β-strand 4 |
| | L489C | β-strand 5 |
| | T511C | β-strand 6 |
| | L529C | β-strand 7 |
| | F572C | β-strand 8 |
| | L595C | β-strand 9 |
| | L613C | β-strand 10 |
| | W635C | β-strand 11 |
| | A714C | β-strand 12 |

| | |
|---|---|
| W739C | β-strand 13 |
| I772C | β-strand 14 |
| F785C | β-strand 15 |
| F804C | β-strand 16 |
| E435C | extracellular loop 1 |
| D464C | extracellular loop 2 |
| D500C | extracellular loop 3 |
| R547C | extracellular loop 4 |
| S604C | extracellular loop 5 |
| E650C | extracellular loop 6 |
| Q664C | extracellular loop 6 |
| Y674C | extracellular loop 6 |
| D685C | extracellular loop 6 |
| D694C | extracellular loop 6 |
| D704C | extracellular loop 6 |
| S755C | extracellular loop 7 |
| D795C | extracellular loop 8 |
